# OEP24.1 involved in carbon allocation is a receptor of piecemeal plastid autophagy in Arabidopsis

**DOI:** 10.64898/2026.04.02.715788

**Authors:** Louis Lambret, Rozenn Le Hir, Jie Luo, Fabien Chardon, Anne Margagne, Céline Masclaux-Daubresse

**Affiliations:** Université Paris-Saclay, INRAE, AgroParisTech, Institute Jean-Pierre Bourgin for Plant Sciences, 78000, Versailles, France

**Author notes:** authors contributed equally (Louis Lambret, Anne Margagne, Céline Masclaux-Daubresse). Jie Luo: College of Horticulture and Forestry Sciences, Huazhong Agricultural University, Wuhan, 430070, China. The author(s) responsible for distribution of materials integral to the findings presented in this article in accordance with the policy described in the Instructions for Authors (https://academic.oup.com/plcell/pages/General-Instructions) is (are): Marmagne A. and Masclaux-Daubresse C. Authors (;).

## Abstract

Macroautophagy is a conserved intracellular catabolic process in eukaryotes that participates in chloroplast degradation, through the selective breakdown of chloroplast components. Selective autophagy of membrane-bound organelles typically requires receptors that bridge organelle membranes and pre-autophagosomal structures. Here we identify OEP24.1 as a new receptor in the selective chloroplast piecemeal autophagy, supporting the degradation of stromal proteins. We found that the β-barrel protein OEP24.1 is located at the outer membrane of plastid envelopes and on bodies budding off plastids into the cytosol and containing stroma proteins. OEP24.1 interacts physically with ATG8 autophagy proteins in a UIM dependent manner. OEP24.1-GFP and RFP-ATG8 colocalize with in mobile autophagosome-like puncta in the cytosol and in autophagic bodies within the vacuole. Delivery of OEP24.1 to vacuole lumen is dependent on active autophagy. OEP24.1 controls carbon allocation at the whole plant level, carbon concentrations in flowering stems and xylem composition. These phenotypes can be explained by the role of OEP24.1 in metabolite diffusion across the chloroplast envelope, and by its involvement in the facilitation of chloroplast quality control through piecemeal autophagy.

## INTRODUCTION

Macroautophagy is an intracellular vesicular process in eukaryotic cells that involves the formation of a double-membrane vesicle, called the autophagosome, which sequesters harmful or unwanted cytoplasmic components ^1^. Through cellular cleansing, macroautophagy maintains homeostasis, organelle quality control, and longevity. The macroautophagy machinery involves numerous proteins (ATG proteins) essential for the formation of the autophagosome, based on the emergence, growth, and closure of the phagophore around autophagic cargoes, as well as the trafficking of the mature autophagosome to the lytic vacuole. The high selectivity of macroautophagy for cargoes depends on the direct or indirect interactions between cargoes and ATG8, a ubiquitin-like protein that is conjugated to lipids and anchored to the membrane of the phagophore upon activation of autophagy. During selective autophagy, e.g. organelle piecemeal autophagy, specific receptors act as intermediates linking cargoes with ATG8 proteins. Ultimately, receptors and cargoes sequestered in autophagosomes are delivered to the lytic compartments for degradation and recycling. The sequestration of receptors or cargoes is supported by the presence of two conserved motifs: the ATG8-interacting motif (AIM) and the ubiquitin-interacting motif (UIM) ^2^. To date, the cargo receptors mediating autophagic organelle recycling have remained largely elusive in plants ^3^.

Chloroplast turnover and quality control are essential for cell longevity, as they preserve photosynthetic capacity and prevent oxidative damage. Chloroplasts are major nitrogen reservoirs in mesophyll cells. Chloroplast decay in response to stress or during developmental leaf senescence is crucial for nutrient use efficiency and resource allocation at the whole-plant level ^4^. However, the coordination and role of autophagy and proteases in chloroplast dismantling remain under debate. Early studies suggested that the chemical fragmentation of stromal proteins by the release of reactive oxygen species (ROS) under stress, through photochemical exchanges, may precede chloroplast decay ^5^. Furthermore, studies have provided evidence indicating that stromal proteins, either native or partially fragmented, could be degraded inside the vacuole ^6–9^. Evidence supporting the role of autophagy in the trafficking of chloroplast components to the vacuole has been provided by several studies, particularly using microscopy ^10–15^.

Several types of vesicles involved in piecemeal chloroplast degradation or trafficking of chloroplast material to the vacuole have been described. Rubisco Containing Bodies (RCBs), which bud at the tips of stromules, are thought to specialize in the export of stroma material. The delivery of RCBs to the vacuole has been shown to depend on the ATG5 autophagy core machinery ^16–21^. However, how RCBs are recognized by the autophagy machinery and sequestered by autophagosomes remains unclear ^16^. The ATI1-PS (ATG8-interacting protein 1-Plastid) bodies are vesicles formed within plastids ^10^. They are likely delivered to the cytosol in an autophagy-independent manner, although the ATI1 protein was shown to interact with ATG8. The budding and detachment of ATI1-PS bodies from the chloroplast envelope do not require the autophagy core machinery. ATI1-PS bodies are plant-specific, and the ATI1 protein was also shown to localize at the endoplasmic reticulum (ER), where it is involved in the starvation-induced reticulophagy pathway, acting as a receptor for MSBP1/MAPR5 cargos ^22,23,10^. It has been shown that ATI1 can bind both stromal and membrane-bound chloroplast proteins, suggesting that the cargoes of ATI1-PS bodies differ from those of RCBs ^10^. Senescence-associated vacuoles (SAVs) are another type of vesicle potentially involved in chloroplast degradation. SAVs are characterized by intense proteolytic activity. The overexpression of SAG12-GFP fusions in chloroplast-containing leaf cells facilitated their observation with confocal microscopy. SAVs have been shown to contain several chloroplast proteins, supporting their role in chloroplast breakdown ^24^ The biogenesis of SAVs, like that of RCBs and ATI1-PS, is stimulated during senescence and by abiotic stresses ^6,16^. Despite the strong induction of SAG12 (Senescence-associated gene 12) and the substantial increase in SAV numbers during leaf senescence, *sag12* mutants only displayed subtle phenotypes and the role of SAVs in chloroplast decay thus remains unclear ^25,26^. CCV vesicles, namely vesicles containing chloroplast vesiculation (CV) proteins, have been shown to contain thylakoid proteins ^15^. Their formation is enhanced by senescence and stress in Arabidopsis. Microscopy showed that CCVs are attached to the envelope of disassembled chloroplasts. Silencing the gene encoding CV protein has been shown to delay chloroplast turnover, while overexpression of CV proteins accelerates it. The trafficking of CCVs to the vacuole is likely independent of autophagy ^15^.

In addition to the piecemeal chloroplast degradation that involves the trafficking of chloroplast material inside vesicles, the degradation of entire chloroplasts has been demonstrated after chloroplasts were shrunken and damaged by either UV-B or high light treatments. Such shrunken chloroplasts have been shown to be engulfed inside the vacuole due to tonoplast invagination processes, a form of micro-autophagy ^27,28^. Whether micro-autophagic chloroplast degradation depends on the canonical ATG machinery still needs further investigation.

Based on comparative proteomics, we identified a chloroplast envelope protein that systematically accumulates in the leaves of autophagy mutants under stress conditions ^29^. This protein, encoded by the At5g42960 locus, is homologous to a pore protein previously isolated from *Pisum sativum* leaves ^30^. By analogy to pea, we named this protein OEP24.1, and we provide here several pieces of evidence suggesting that OEP24.1 serves as a receptor for chloroplast piecemeal autophagy ^31^.

## RESULTS

### The OEP24.1 protein accumulates in leaves of autophagy mutants due to impaired autophagy activity

Proteomic analyses performed on autophagy mutants by Havé et al. (2019) enabled the identification of proteins that abnormally accumulate in autophagy-deficient plants under stress conditions ^29^. In their study, authors compared plants carrying *atg5* mutations in two different genetic backgrounds (Col-0 and *sid2*) and under three growth conditions (nitrate- and sulfate-sufficient supply, nitrate limitation, and sulfate limitation) in order to identify *atg5*-dependent protein overaccumulation. Among the large set of proteins accumulating in *atg5* mutants, we focused on the OEP24.1 protein, predicted to localize to the outer membrane of the chloroplast envelope. OEP24.1 accumulated in the leaves of 60-day-old *atg5* rosettes regardless of stress conditions or salicylic acid synthesis (Fig. 1A). In a second experiment consisting of dark-stress treatment on individual leaves (unpublished), OEP24.1 was still found to be overabundant in *atg5* leaves (Fig. 1B).

**Figure 1:**
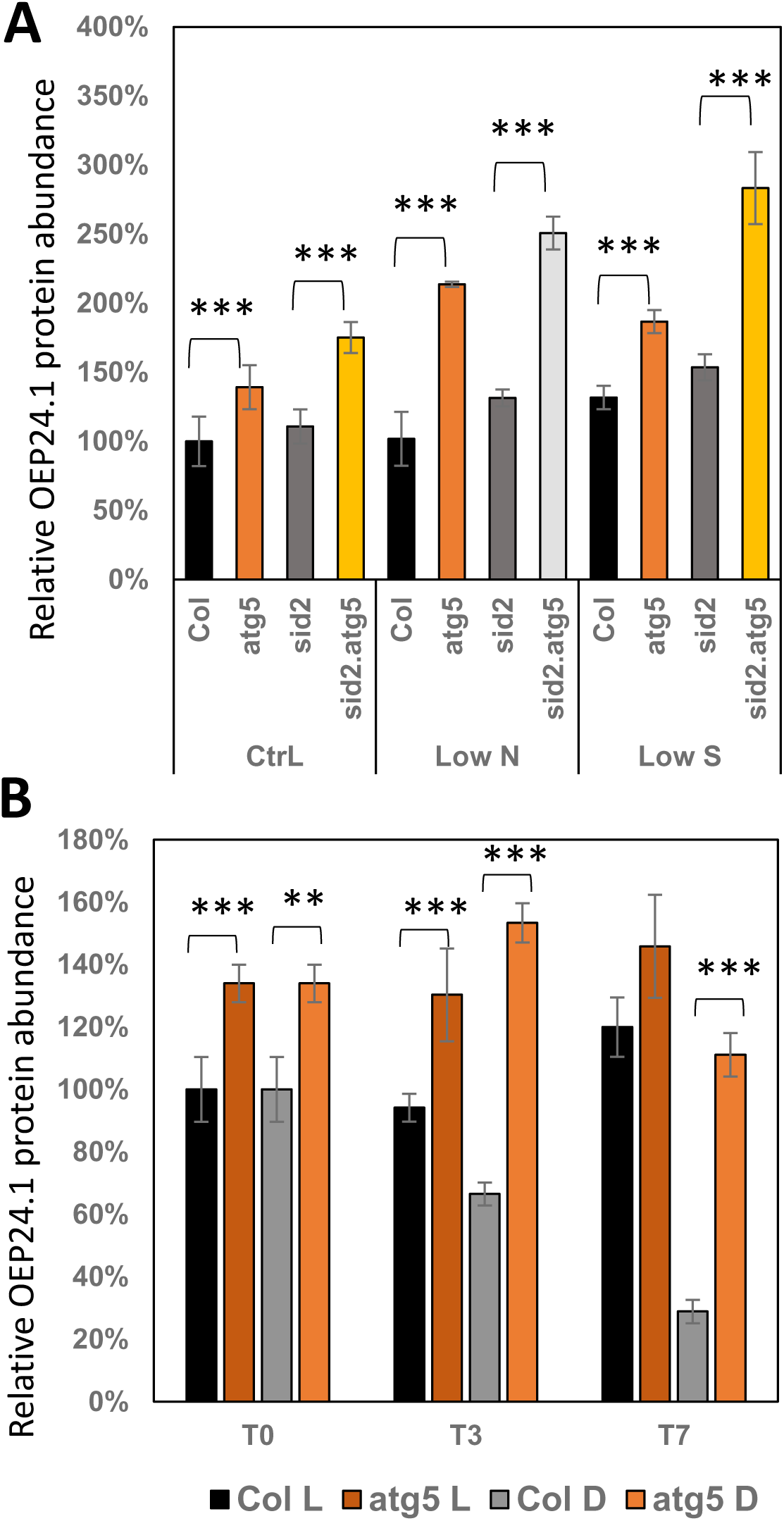
OEP24.1 protein over accumulates in autophagy mutants under N, S, and C starvation conditions. (A) The relative abundances of the OEP24.1 protein (At5g42960) in rosettes of wild-type (Col) and *atg5*, *sid2*, and *sid2 atg5* mutants were calculated from proteomic data obtained by Havé *et al.* (2019). Plants were grown under control (Ctrl), low-nitrate (Low N), and low-sulfate (Low S) conditions. Relative abundances were normalized to the values obtained for wild-type (Col) plants under control conditions. (B) Relative abundances of the OEP24.1 protein in rosettes of wild-type (Col) and *atg5* plants were extracted from unpublished proteomic data (Havé, Lambret *et al.*, unpublished). Proteomic analyses were performed on individual leaves either subjected (D) or not subjected (L) to dark stress. T0, T3, and T7 indicate the number of days of treatment. Means ± SD of three independent replicates are shown. Significant differences between genotypes are indicated (ttests; **P < 0.01; ***P < 0.001).

Since nutrient stresses trigger chloroplast decay and since chloroplasts can be partially degraded through autophagy ^32^, we examined OEP24.1 degradation upon autophagy induction *via* a GFP-cleavage assay using *Arabidopsis* OEP24.1-GFP transformants (Fig. 2). This assay relies on the relative stability of free GFP in the vacuole, such that autophagy-dependent delivery of GFP-tagged proteins to the vacuole results in the accumulation of cleaved GFP detectable by immunoblotting. Seedlings grown *in vitro* on agar plates were transferred to nutrient rich liquid medium (+NC) or to liquid medium lacking both nitrogen and carbon sources (-NC). In wild-type seedlings, free GFP was detected in the -NC samples but not in the +NC samples, indicating that OEP24.1-GFP was not cleaved when autophagy was not induced (+NC), but underwent degradation upon autophagy activation (-NC) (Fig. 2A). In contrast, no free GFP was released from OEP24.1-GFP degradation in *atg5* seedlings under either -NC or +NC conditions (Fig. 2B), indicating that OEP24.1 degradation is autophagy-dependent. Treatment of wild-type seedlings with concanamycin A in the in autophagy-inducing buffer (-NC + CA) prevented OEP24.1-GFP cleavage, demonstrating that OEP24.1-GFP degradation during autophagy occurs within the vacuole (Fig. 2A). Consistently, free GFP was also absent in *atg5* seedlings transferred to -NC +CA (Fig. 2B)

**Figure 2:**
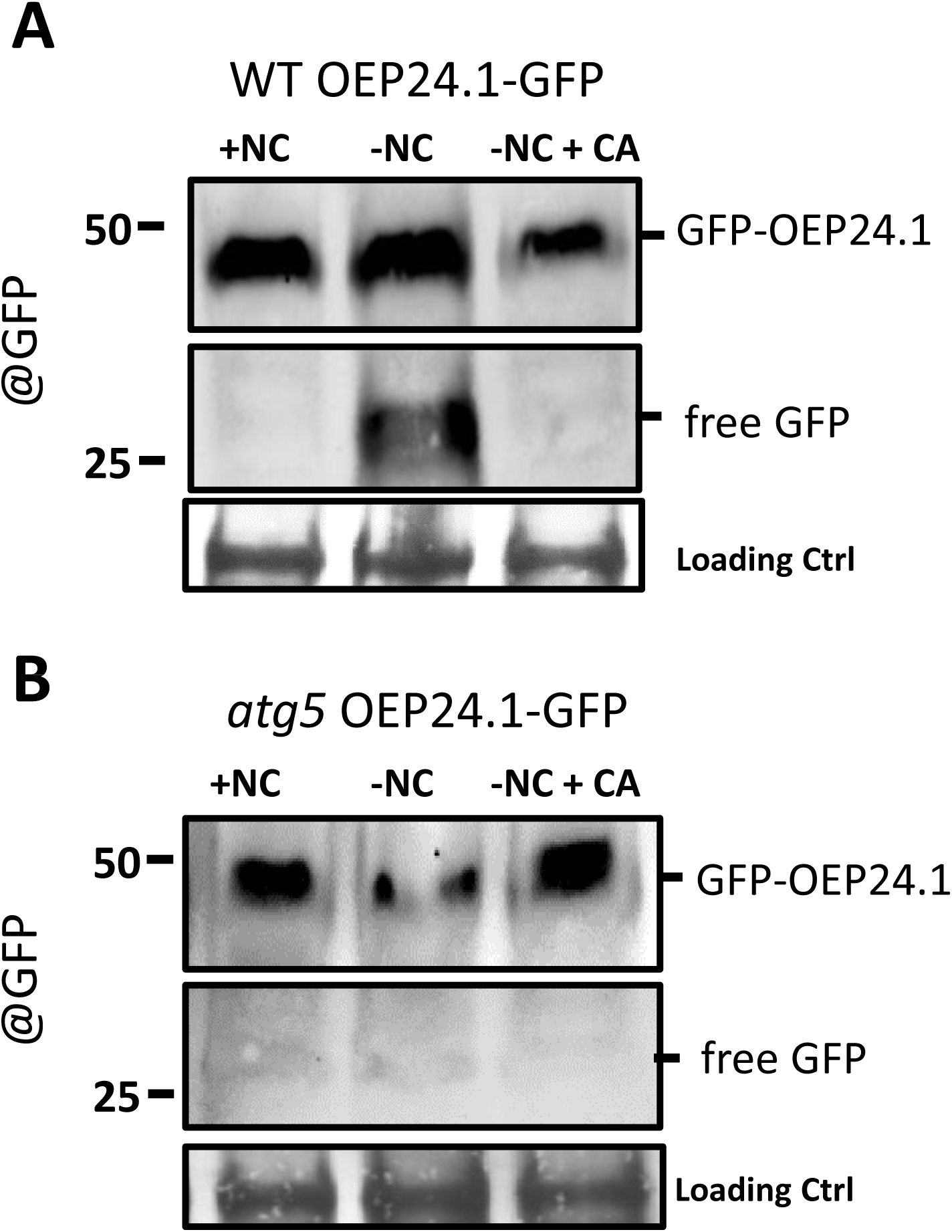
Degradation of OEP24.1–GFP depends on autophagic flux. The degradation of the OEP24.1–GFP fusion protein and the release of free GFP into the vacuole were investigated in WT and *atg5* seedlings expressing the OEP24.1–GFP construct. Ten-day-old OEP24.1–GFP seedlings grown were transferred from nutrient-rich (MS/2) agar plates to either nutrient-rich liquid medium (+NC), nutrient-starvation medium (−NC; 3 mM MES, pH 5.7), or nutrient-starvation medium supplemented with concanamycin A (1 μM; −NC + CA) for 4 hours. Immunoblot analyses of OEP24.1–GFP and free GFP levels in seedling extracts from WT (A) and *atg5* mutants (B) were performed using commercial anti-GFP antibodies. Protein loading was controlled using amido black staining.

### Alphafold predictions involving UIM and crumble AIM motifs in OEP24.1

We first generated an AlphaFold3 model of OEP24.1 to assess its folding and potential orientation at the chloroplast outer membrane. The predicted structure displayed the canonical porin-like fold with a 12 β-barrel domain, in agreement with previous descriptions of OEP24 proteins (Fig.3A; ^33^). *In silico* annotation highlighted (i) the GSFXIV putative AIM-like sequence (Fig.3AB in blue), hereafter referred to as a crumble-AIM, since it is only retrieved upon insertion of an additional residue in the X-LIR search algorithm, and (ii) two putative UIM motifs (Fig.3AB, in green). To further evaluate the ability of OEP24.1 to be embedded into a lipid bilayer, we modelled OEP24.1 in the presence of >50 lipid ligands (Fig.3C). The protein spontaneously embedded into the bilayer, consistently forming a stable β-barrel spanning the membrane (Fig.3C), with both the N and C terminals at the surface of the membrane at the cytosol side (Fig.3D). Then, AlphaFold3 multimer predictions were generated between OEP24.1 and each of the nine Arabidopsis ATG8 proteins (ATG8a–i), (Supplemental Fig.S1). For ATG8a–g, the C-terminal residues corresponding to the downstream sequence from the ATG4 cleavage site, were removed to mimic the lipidated forms of these ATG8, while ATG8h and ATG8i were modelled in their native sequence. In all cases, predictions yielded low ipTM scores (0.1 to 0.29) and poorly defined PAE maps (Supplemental Fig.S1). In several cases, including ATG8b, ATG8c, ATG8d and ATG8i, ATG8 protein appeared to insert into the OEP24.1 β-barrel, thereby perturbing the pore architecture but lacking any interaction with the predicted AIM or UIMs. In contrast, the predicted interaction with ATG8e was localized to the cytosolic region of OEP24.1, which corresponds to the expected binding mode for ATG8, but still has a very low iPTM score (0.1). Although AlphaFold3 failed to reveal any robust interaction, we hypothesized that additional regulatory features (such as post-translational modifications, membrane context, or conformational rearrangements) may be required for OEP24.1-ATG8 recognition *in vivo*.

**Figure 3:**
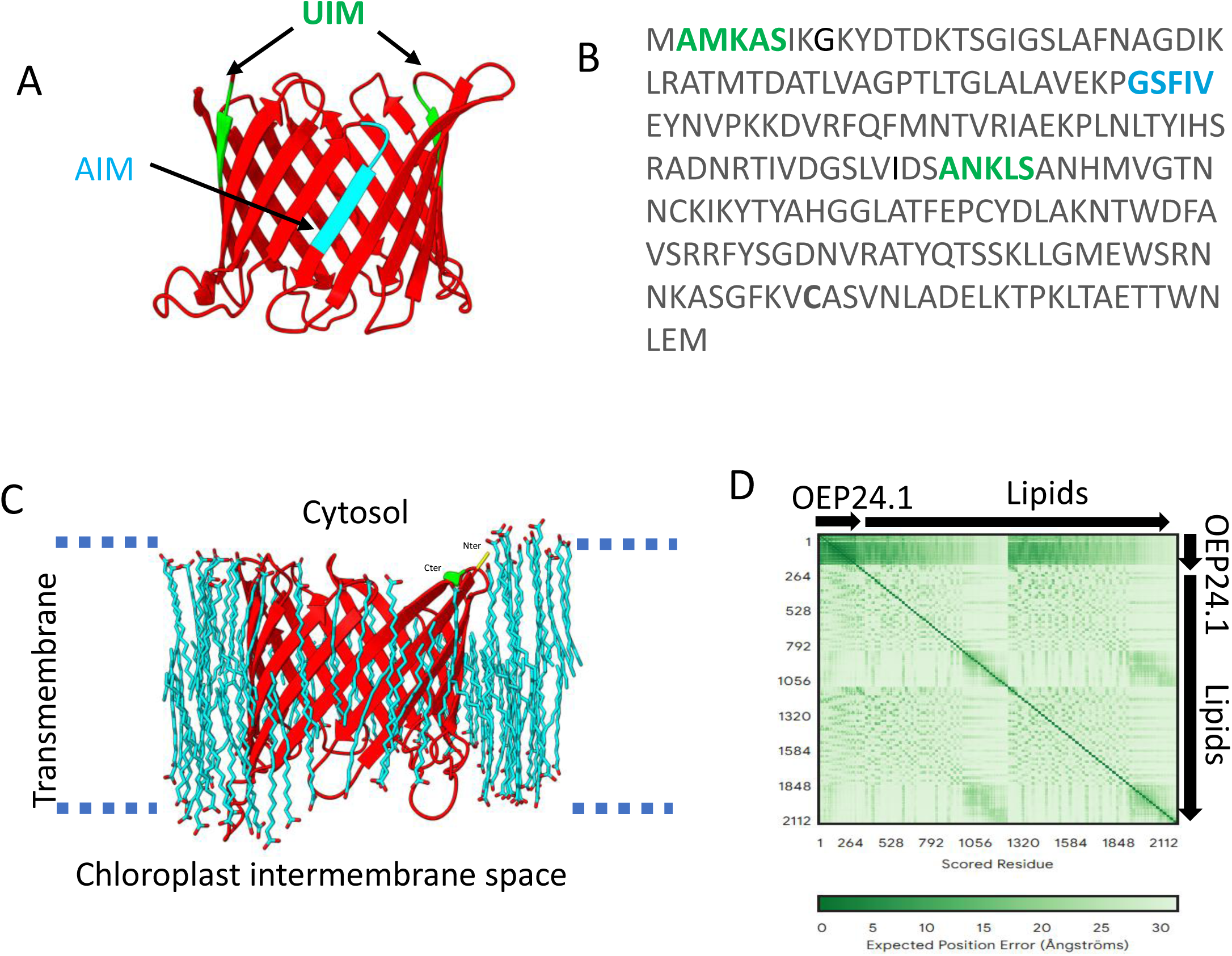
AlphaFold models and structural features of OEP24.1. (A) Predicted structure of OEP24.1 obtained using AlphaFold3, showing a β-barrel porin fold. One putative ATG8-interacting motif (AIM; cyan) and two ubiquitin-interacting motifs (UIMs; light green) are highlighted. (B) Amino acid sequence of OEP24.1 with the AIM (cyan) and UIMs (light green) highlighted. (C) AlphaFold3 model of OEP24.1 embedded in a lipid environment containing 50 lipids to illustrate membrane insertion. The protein is oriented with its C-terminus (green) and N-terminus (yellow) exposed to the cytosol, while the β-barrel spans the chloroplast outer envelope membrane. (D) Predicted aligned error (PAE) plot corresponding to the model shown in panel (C).

### The OEP24.1 protein binds ATG8 isoforms at UDS site

To experimentally assess OEP24.1–ATG8 interactions beyond *in silico* predictions, we tested the physical interaction of OEP24.1 with the nine ATG8 isoforms using three different *in vivo* approaches. Yeast two hybrid (Y2H) pairwise interaction assays performed with the nine ATG8a-i isoforms (Fig.4A; Supplemental Fig. S2) showed that all ATG8 isoforms interacted with OEP24.1, except ATG8a. We then used ATG8 isoforms carrying alanine substitutions in the LDS (Y_51_H_52_ -> A_51_A_52_) or the UDS (I_78_F_79_I_80_ -> A_78_A_79_A_80_) sites to determine whether the OEP24.1-ATG8 interactions depended on the AIM or the UIM motifs (Fig.4B). Considering the AlphaFold3 models, we chose to modify the ATG8e and ATG8d isoforms to test Y2H interactions with OEP24.1. In the case of ATG8d, the interaction with OEP24.1 is predicted to occur within the lipid bilayer, whereas for ATG8e, it is predicted to occur at the cytosolic surface of the membrane. While the ΔLDS variants of ATG8d andATG8e still interacted successfully with OEP24.1, their interactions with ΔUDS variants was weak (Fig.4B), demonstrating that binding of OEP24.1 to ATG8d or ATG8e mainly depends on UIM-UDS interactions. It was surprising to observe that interactions between OEP24.1 and the ΔLDS ATG8e and ΔLDS ATG8d variants was stronger than interactions of OEP24.1 with the native ATG8e and ATG8d. The presence of LDS likely results in a lower accessibility of the UDS site to OEP24.1.

**Figure 4:**
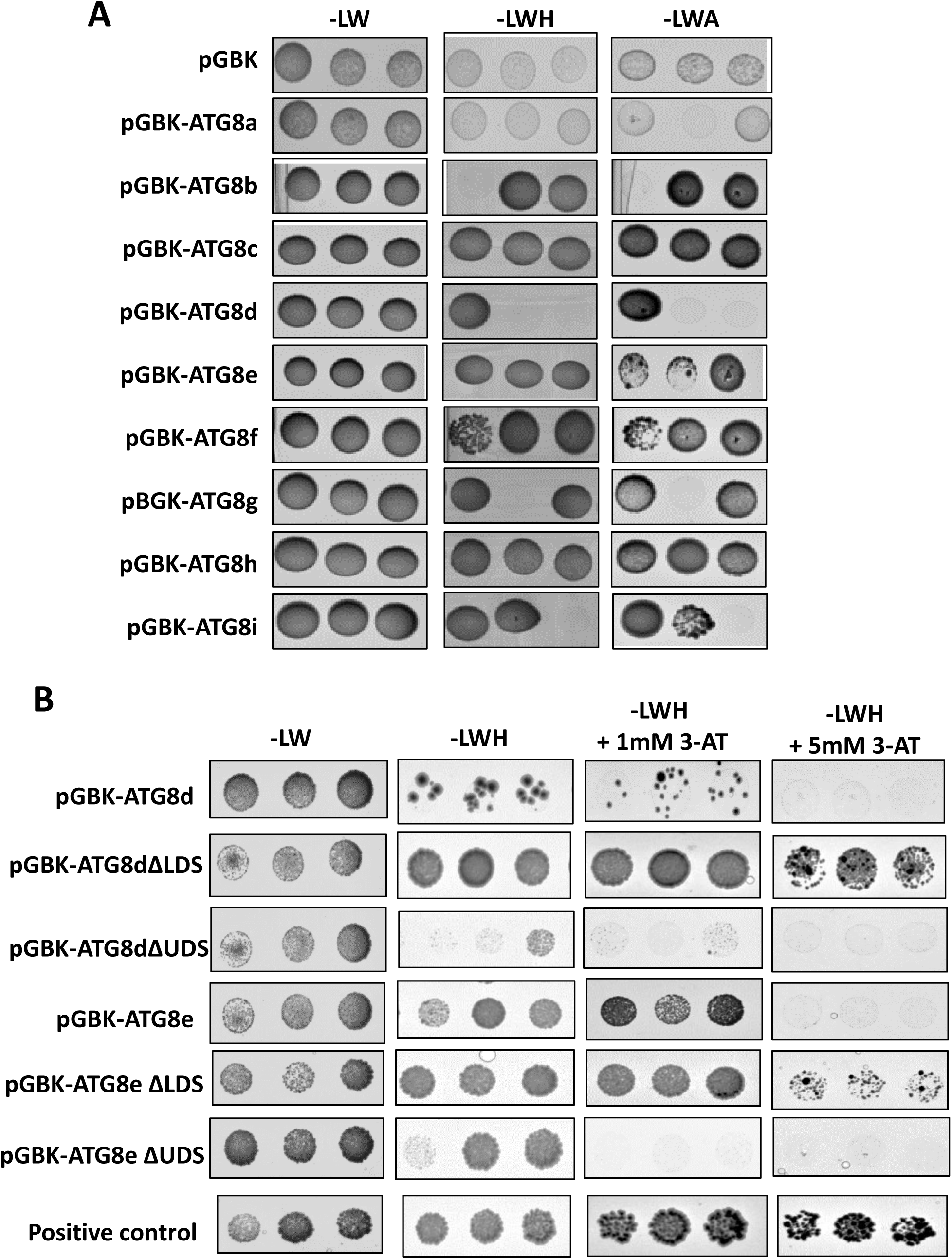
OEP24.1 interacts with ATG8 isoforms in a UIM–UDS-dependent manner. ATG8a–i isoforms were used as baits (BD fusions), and OEP24.1 was used as prey (AD fusion). (A) Wild-type ATG8a–i isoforms used as baits. Yeast cells harboring both bait and prey expression vectors were grown on medium lacking histidine (−LWH) or adenine (−LWA). (B) Mutated ΔUDS and ΔLDS variants of ATG8e and ATG8d were used as baits. Yeast cells harboring both bait and prey expression vectors were grown on medium lacking histidine (−LWH) or on −LWH medium supplemented with increasing concentrations of 3-AT to assess interactions between partners. In (A) and (B), results show the growth of three representative clones after yeast mating. Experiments were repeated at least three times (n ≥ 3) with similar results. The positive control consisted of the interaction between ATG8g (BD) and NBR1 (AD). The negative control consisted of testing the interaction between empty pGBK (BD) and OEP24.1 (AD).

To test OEP24.1–ATG8 interactions *in planta*, we performed bimolecular fluorescence complementation (BiFC) assays, which allow visualization of protein–protein interactions through the reconstitution of a functional YFP fluorophore. Co-expression of nYFP–OEP24.1 with cYFP-tagged ATG8 isoforms in *Nicotiana benthamiana* leaves resulted in a clear YFP signal for the three isoforms tested (ATG8d, ATG8g and ATG8i), indicating a physical interaction *in planta* (Fig.5A; Supplemental Fig.S3).

**Figure 5:**
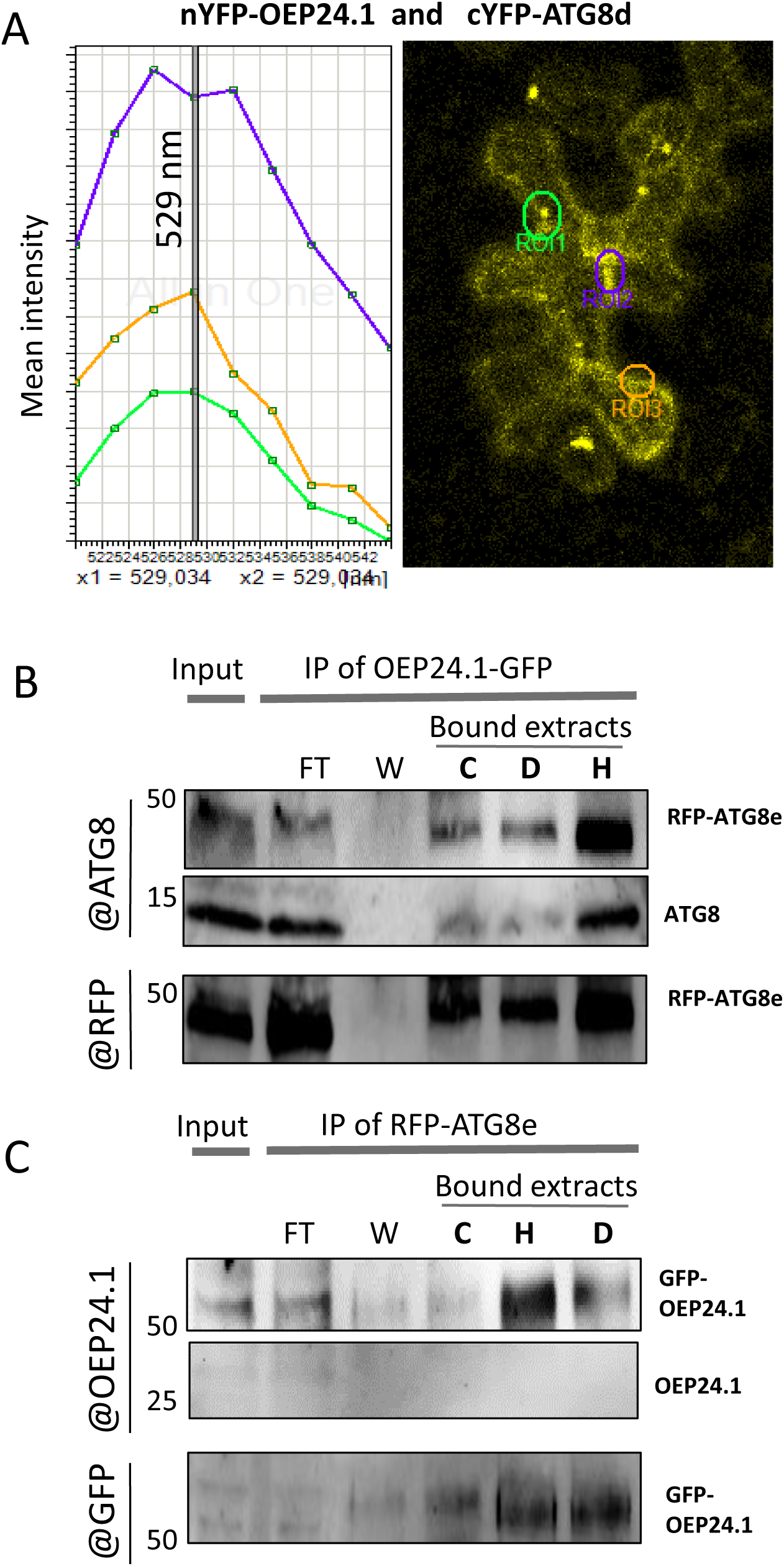
ATG8 and OEP24.1 interact *in planta*. (A) Bimolecular fluorescence complementation (BiFC) was performed by co-agroinfiltrating *Nicotiana benthamiana* leaves with the pUbi::nYFP-OEP24.1 and pUbi::cYFP-ATG8d constructs in the presence of P19. Confocal imaging was performed two days after leaf infiltration. Excitation of leaf tissues at 514 nm induced fluorescence signals with an emission peak at 529 nm, corresponding to YFP. Regions of interest (ROIs) used for spectral analyses are indicated by circles; the colors of the ROI circles correspond to the colors of the spectra. Other interactions tested with ATG8g and ATG8i, as well as controls, are shown in Supplemental Fig.S3. (B, C) Immunoprecipitation assays were performed using ten-day-old *Arabidopsis* transgenic lines expressing OEP24.1–GFP and RFP–ATG8e fusion proteins. Seedlings were grown on ½ MS medium for 10 days and then either maintained under control conditions (lane C), transferred to darkness (lane D), or subjected to heat stress (lane H). Only the input, flow-through, and wash fractions for the control condition are shown. Similar signal intensities were obtained for the input, flow-through, and wash fractions under dark and heat treatments (see Supplemental Fig.S4). Immunoprecipitation experiments were repeated three times (n = 3) with similar results. FT, flowthrough; W, wash elution.

Co-immunoprecipitation was then performed using Arabidopsis seedling carrying both RFP-ATG8e and the OEP24.1-GFP fusion proteins. Autophagy was either activated or not by applying dark or heat stress treatments. Immunoprecipitation of OEP24.1-GFP resulted in the recovery of both endogenous ATG8 proteins and the RFP-ATG8e fusion protein, as revelated by immunoblotting using anti-ATG8 and anti-RFP antibodies, respectively, in dark-stressed and heat-stressed seedings (Fig.5B). Immunoprecipitation performed on *p35S::GFP* control plants confirmed that ATG8 binds specifically to OEP24.1 and not to GFP alone (supplemental Fig.S4A). Conversely, in immunoprecipitation targeting RFP-ATG8, GFP-OEP24.1 was detected in the bound fractions by immunoblotting with anti-GFP antibodies in seedlings grown under control conditions, with increased recovery in dark-stressed seedlings and the highest recovery observed under heat-stress (Fig.5C). Using the dz297 antibody raised against OEP24.1, whose specificity was validated comparing extracts from wild-type and *oep24.1* CRISPR mutant plants (Supplemental Fig.S5), endogenous OEP24.1 could not be detected in the crude extract, the flow through fraction, or the bound fraction from RFP-ATG8e immunoprecipitation (Fig.5C). This lack of detection is likely due to the low expression of the *OEP24.1* gene at seedling stage. Nevertheless, the OEP24.1-GFP fusion protein was readily detected using dz297 antibody. Co-immunoprecipitation assays performed in *Nicotiana benthamiana* leaves transiently expressing OEP24.1-GFP alone, together with GFP-ATG8b, or with RFP-ATG8d further confirmed the interaction between OEP24.1 and these ATG8 isoforms (Supplemental Fig.S4).

Altogether, these experiments demonstrate that OEP24.1 interacts with several ATG8 isoforms in yeast and *in planta*, binding likely relying on the UDS of ATG8.

### OEP24.1 bodies bud off from the chloroplast and etioplast envelopes, contain stromal proteins, and are transferred to the vacuolar lumen in a process that requires active autophagy

To determine the subcellular localization of OEP24.1 in leaf cells and whether its interaction with ATG8 leads to sequestration within autophagosomes and autophagic bodies, OEP24.1 fusion proteins were expressed transiently in *Nicotiana benthamiana* leaves and stably in *Arabidopsis* plants. In both *Arabidopsis* leaves (Fig. 6) and *Nicotiana benthamiana* agroinfiltrated leaves (Supplemental Fig. S6), OEP24.1-GFP localized to chloroplasts. Fluorescence was observed in large chloroplasts (∼5 μm), mainly in mesophyll cells, and in small chloroplasts (∼1 μm) with lower chlorophyll autofluorescence and predominantly present in epidermal cells. Although the precise localization of OEP24.1-GFP within large chloroplasts of mesophyll cells was difficult to determine (Fig. 6), the ring-shaped fluorescence surrounding the small chloroplasts of epidermis indicated localization at the chloroplast envelope (Fig. 6). Consistently, OEP24.1-GFP also localized at the envelope of root etioplasts. In the *atg5* background, OEP24.1 was observed extending from stromules (Supplemental Fig. S7).

**Figure 6:**
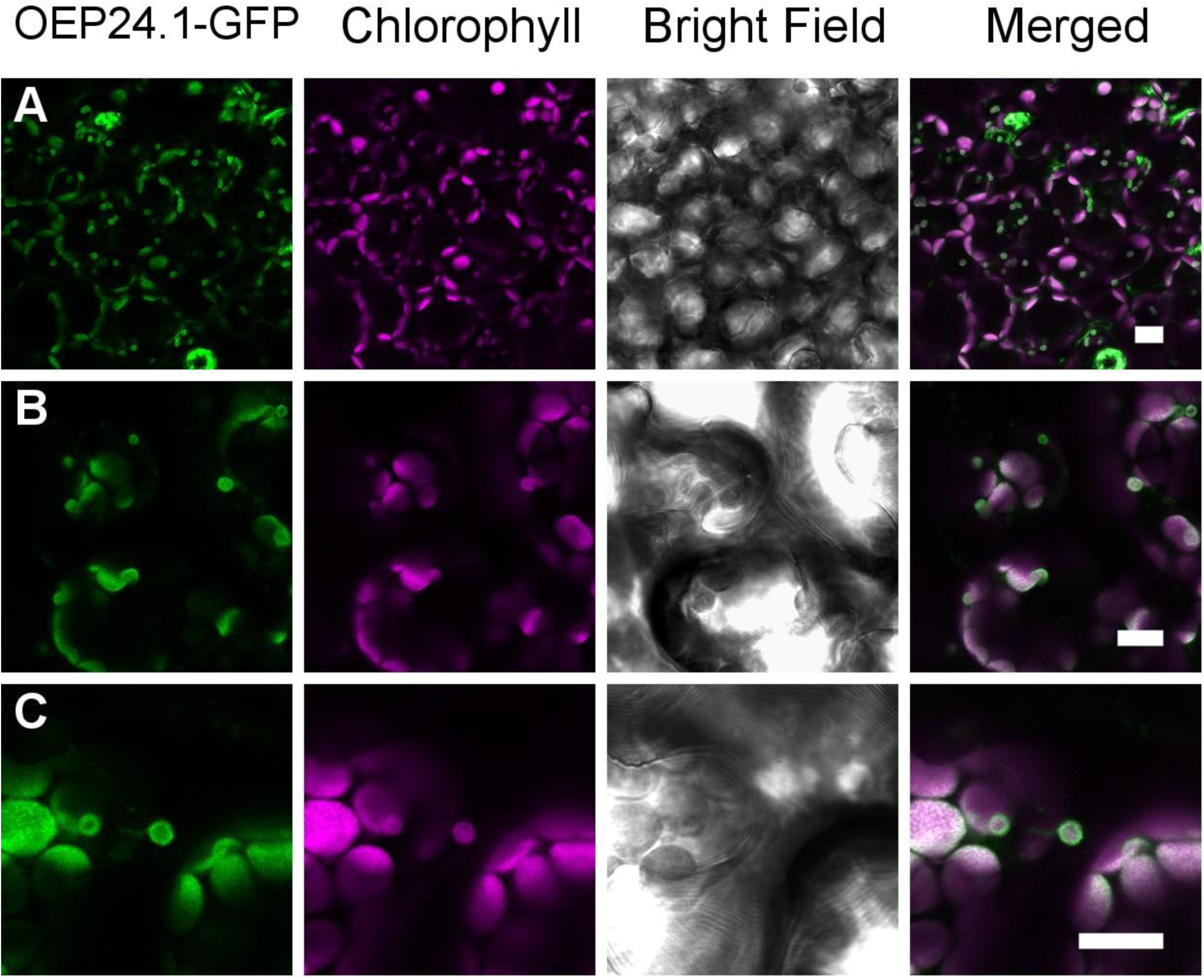
OEP24.1 localizes to the chloroplast envelope in *Arabidopsis* leaf cells. Leaves from 15-day-old wild-type *Arabidopsis* seedlings expressing OEP24.1–GFP were analyzed by confocal microscopy. Three representative optical sections are shown. Green fluorescence corresponds to the OEP24.1–GFP fusion protein, while magenta fluorescence corresponds to chloroplast autofluorescence. Bright-field images show cell boundaries, indicating that small chloroplasts (<1 μm) are most likely located in epidermal cells, whereas larger chloroplasts are found in mesophyll cells. Imaging was repeated more than three times (n > 3), and similar results were observed in agroinfiltrated *Nicotiana benthamiana* leaves (supplemental Fig.S6). Scale bars: 10 μm.

To confirm that OEP24.1-GFP is delivered to the vacuole lumen *via* autophagy, seedlings were treated with Concanamycin A (CA) prior to imaging. In WT cells, GFP fluorescence was readily detected in the vacuole lumen of mesophyll cells (Fig. 7; Movies M1–M4), root cells (Fig.7; Movies M5–M7), and hypocotyls (Supplemental Fig. S8; Movies M8–M10). In contrast, no vacuolar signal was detected in *atg5* cells (Supplemental Fig. S9). Colocalization of RFP-ATG8e and OEP24.1-GFP was then observed in WT seedlings of Arabidopsis upon CA treatment (Fig. 8; Supplemental Fig. S10) and in agroinfiltrated *N. benthamiana* cells (Fig. 9; Supplemental Fig. S11). In both species, RFP and GFP fluorescence signals merged in cytosolic autophagosome-like structures located near chloroplasts (Fig. 8, Fig. 9, Supplemental Fig. S11; Movies M11–M12). In Arabidopsis, colocalization was also observed, inside the vacuole lumen of leaf epidermal and root cells following CA treatment, as mobile aggregates and as autophagic bodies characterized by rapid Brownian motion (Fig. 8; Movies M13–M15). In mesophyll cells, the autophagosomes in which OEP24.1-GFP and RFP-ATG8e colocalized lacked chlorophyll autofluorescence (Fig. 7, Fig. 9; Supplemental Fig. S11; Movie M12), consistent with these structures corresponding to Rubisco-containing bodies ^16^. In order to verify that OEP24.1 bodies contain stroma proteins, we agroinfiltrated *N. benthamiana* leaves with plasmids carrying OEP24.1-GFP and fluorescent marker of stromal protein (pt-yk or pt-ck) ^34^. Using confocal microscopy, we could observe colocalization of OEP24.1-GFP and pt-ck CFP (or pt-yk YFP) signals at chloroplast and stromules (Supplemental Fig. S12) and more importantly on spheric structures budding from chloroplasts (Fig. 10). We also observed spherical structures, still labelled with OEP24.1-GFP and stroma-CFP emerging from chloroplasts within the cytosol (Fig. 11). We could not detect any chlorophyll fluorescence within OEP24.1 bodies, neither at budding site nor when bodies were free and mobile in the cytosol of mesophyll cells (Fig. 10 and 11; Movie M16).

**Figure 7:**
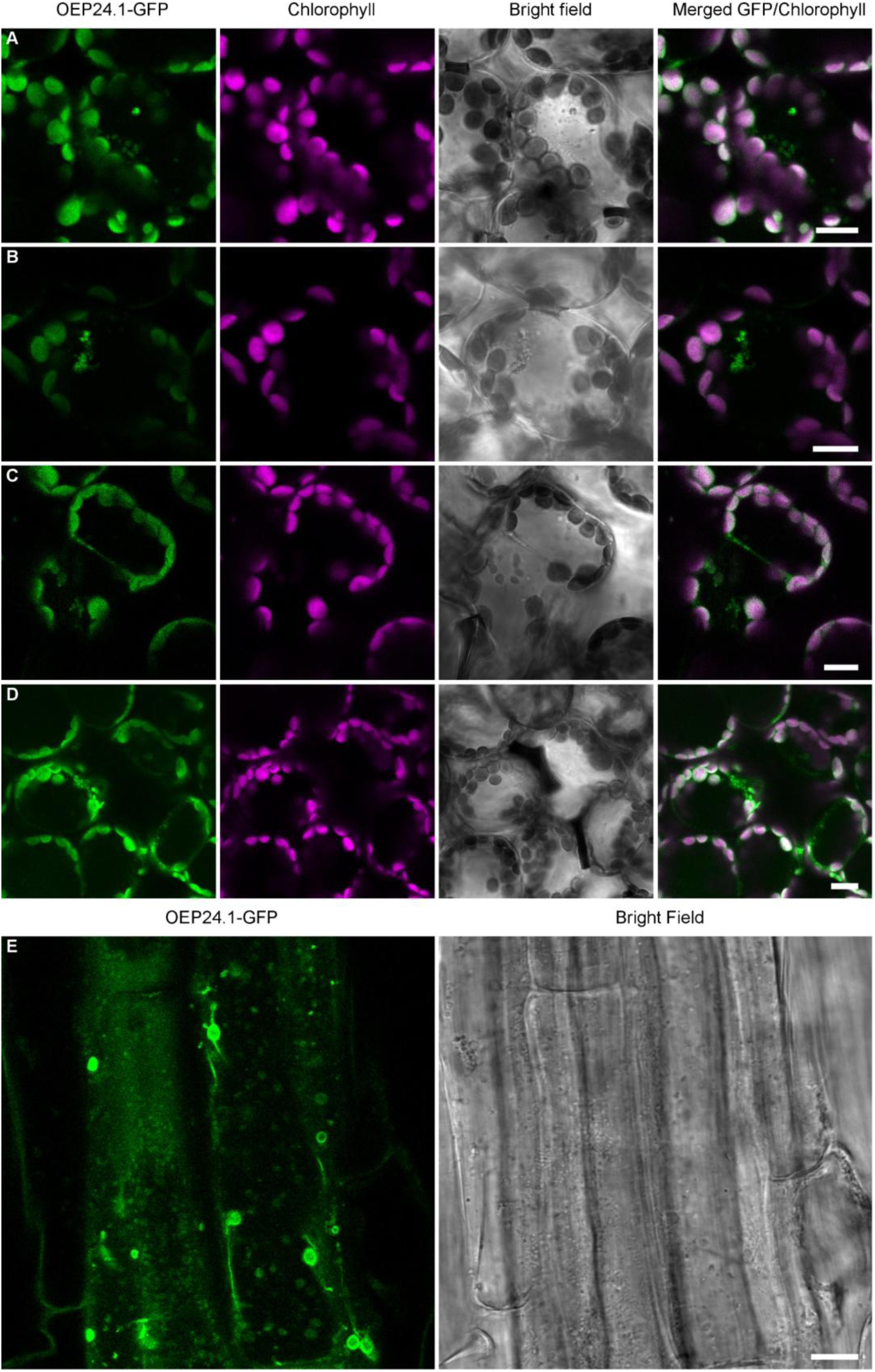
OEP24.1 is transported to the vacuolar lumen of leaf and root cells upon concanamycin A treatment. Leaves and roots of wild-type *Arabidopsis* seedlings expressing OEP24.1-GFP were analyzed by confocal microscopy. Twenty-four hours prior to microscopic observation, plants were treated with 1 μM concanamycin A, an inhibitor of vacuolar H+-ATPase activity and thus of vacuolar proteolytic degradation. As in control conditions (Fig. 6), GFP fluorescence was observed at plastid envelopes and in autophagosomes. Upon concanamycin A treatment, additional localization was detected as aggregates within the vacuole lumen and as mobile autophagic bodies. (A–D) Representative images of mesophyll and epidermal cells. (E) Representative root cell. Imaging was repeated three times (n = 3). Scale bars: 10 μm. Associated movies: M1–M5.

**Figure 8:**
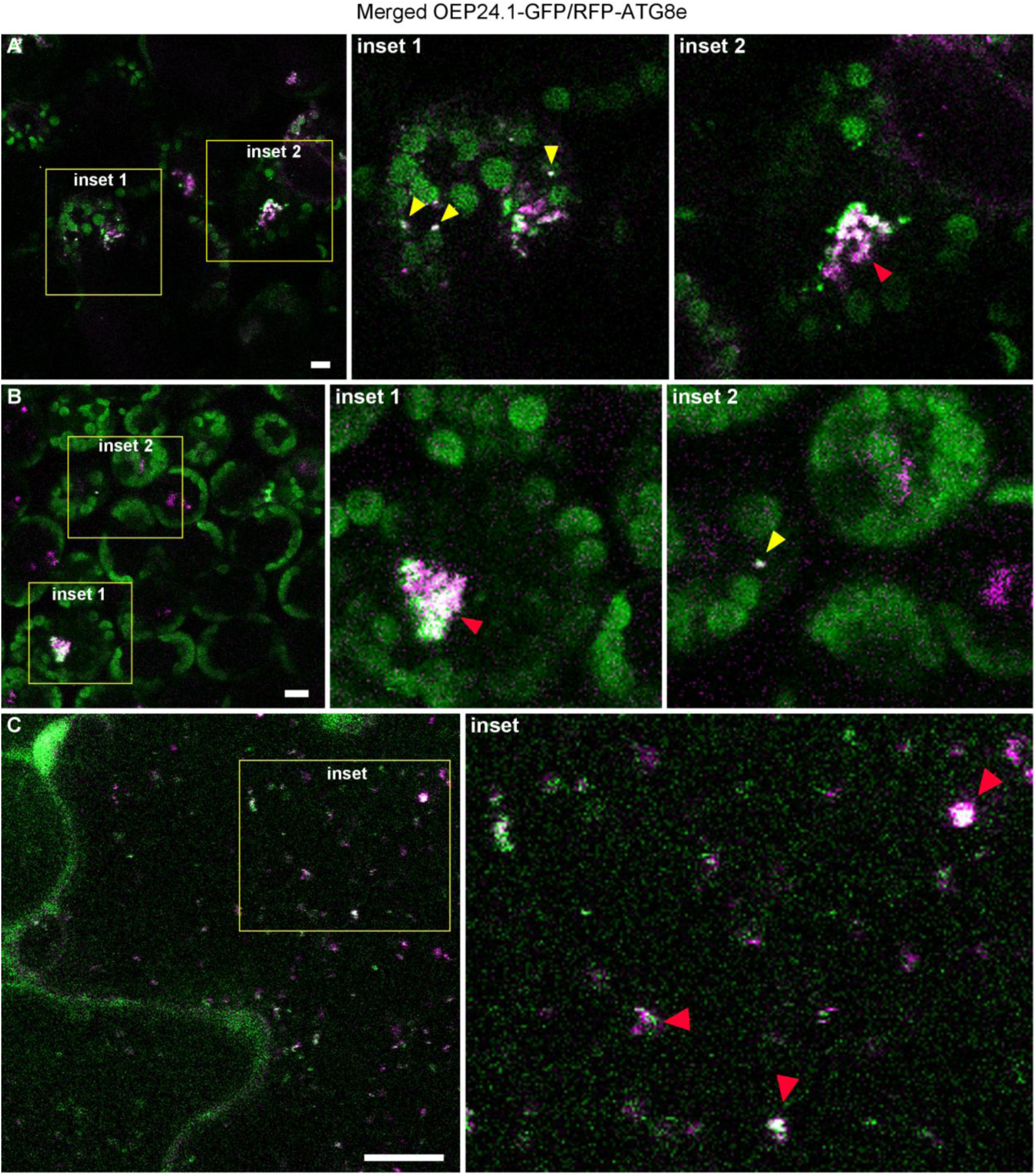
OEP24.1 and ATG8 colocalize in autophagosomes and autophagic body aggregates in the vacuole of wild-type *Arabidopsis* cells upon concanamycin A treatment. Leaves of 15-day-old wild-type *Arabidopsis* seedlings expressing OEP24.1-GFP and RFP-ATG8e were analyzed by confocal microscopy. Seedlings were treated with 1 μM concanamycin A 24 hours prior to imaging. (A, B) Within the vacuole of mesophyll cells, OEP24.1 and ATG8e colocalize in aggregated autophagic bodies (red arrowheads). In the cytosol, colocalizations (yellow arrowheads) correspond to autophagosomes, often located near chloroplasts. (C) In epidermal cells, colocalization is visible in autophagic bodies (red arrowheads) exhibiting Brownian motion (see Movies M13 and M14). Scale bars: 10 μm.

**Fig. 9:**
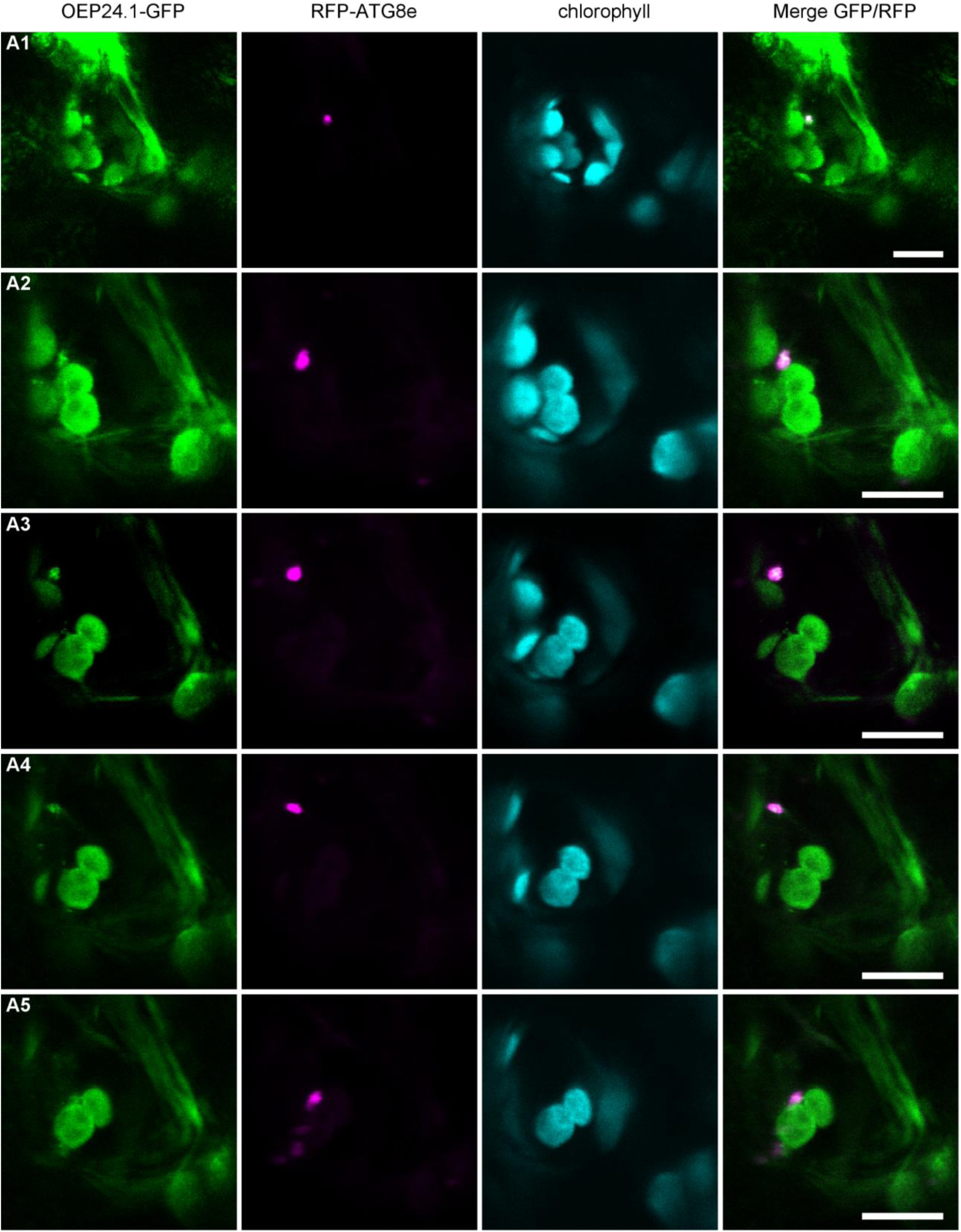
OEP24.1 and ATG8e colocalize in moving autophagosomes in *Nicotiana benthamiana*. Mesophyll cells of *Nicotiana benthamiana* transiently expressing RFP-ATG8e and OEP24.1-GFP constructs following agroinfiltration were analyzed by confocal microscopy. Panels A1–A5 show timelapse images acquired 2 days after agroinfiltration. Green fluorescence corresponds to the OEP24.1–GFP fusion protein, magenta fluorescence corresponds to RFP–ATG8e-labeled autophagosomes, and cyan fluorescence corresponds to chloroplast autofluorescence. The last image in each panel shows a merge of all signals. Scale bars: 10 μm.

**Fig. 10:**
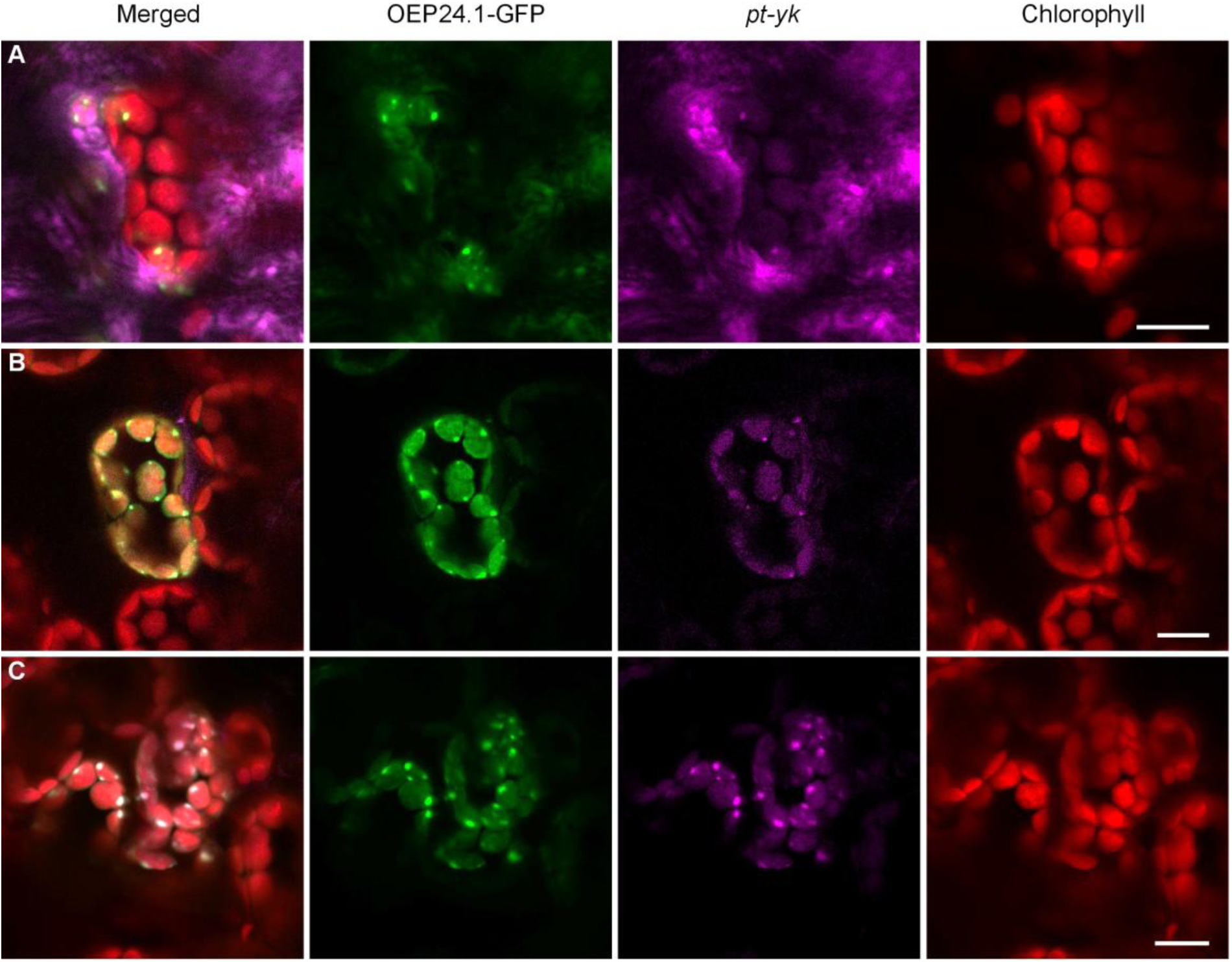
Signals of OEP24.1-GFP and *pt-yk* fusions co-colocalize in puncta located at the surface of chloroplasts in. Mesophyll cells of *Nicotiana benthamiana* transiently expressing and OEP24.1-GFP and *pt-yk* constructs following agroinfiltration were analyzed by confocal microscopy. Panels A-C show images acquired 2 days after agroinfiltration. Green fluorescence corresponds to the OEP24.1–GFP fusion protein, magenta fluorescence corresponds to YFP fused to stromal protein (*pt-yk*), and red fluorescence corresponds to chloroplast autofluorescence. The firstt image in each panel shows a merge of all signals. Scale bars: 10 μm.

**Fig. 11:**
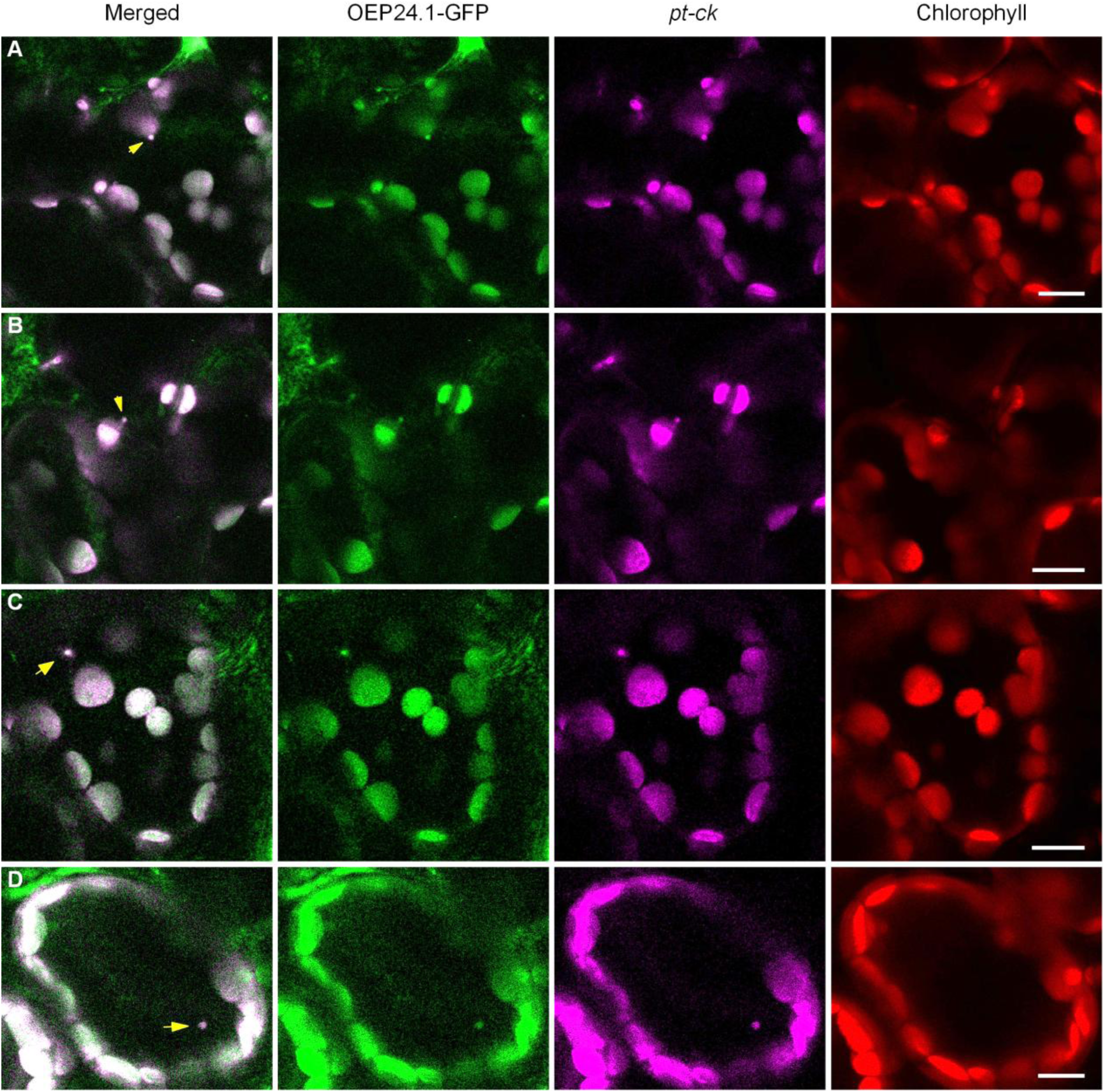
Signals of OEP24.1-GFP and *pt-ck* fusions co-colocalize in puncta located at the surface of chloroplasts (A), detaching from chloroplasts (B) or mobile in cytosol (C,D). Mesophyll cells of *Nicotiana benthamiana* transiently expressing and OEP24.1-GFP and *pt-ck* constructs following agroinfiltration were analyzed by confocal microscopy. Panels A-D show images acquired 2 days after agroinfiltration. Green fluorescence corresponds to the OEP24.1–GFP fusion protein, magenta fluorescence corresponds to CFP fused to stromal protein (*pt-ck*), and red fluorescence corresponds to chloroplast autofluorescence. The first image in each panel shows a merge of green and magenta signals and arrows point at co-localisations. Scale bars: 10 μm.

### OEP24.1 specifically modulates stem carbon allocation and xylem secondary cell wall composition

To investigate the physiological role of OEP24.1, particularly in the selective autophagy and chloroplast turnover during leaf senescence, we generated *OEP24.1* overexpressing lines and CRISPR mutants, aiming to compare their phenotypes with wild type as well as with autophagy mutants and ATG8 overexpressors ^4^. We obtained several independent CRISPR mutants (Supplemental Fig. S13) and three independent *OEP24.1* overexpressing lines (Supplemental Fig. S14). The two CRISPR mutations selected produced truncated proteins that are unable to form functional intramembrane pore channels (Supplemental Fig. S13).

Phenotyping of mutants and overexpressing lines was conducted under both nitrate-sufficient and nitrate-limiting conditions, using the same ^15^N labelling protocol previously employed to reveal phenotypes associated with autophagy stimulation (^35^;^4^) or autophagy deficiency ^36^. Across three independent experiments, each including at least four plant replicates per genotype, no obvious or significant differences were observed in plant development or leaf senescence progression between OEP24.1 mutants, overexpressing lines, and wild type (Supplemental Figs. S14, S15). ^15^N labelling experiments, performed to assess nitrogen and carbon partitioning and nitrogen remobilization - as typically altered in autophagy mutants and *ATG8* overexpressors ^4^ - revealed no significant changes, except for a mild stem phenotype (Supplemental Fig.S15). Under both high- and low-nitrate conditions, the *OEP24.1* CRISPR mutants #4.2 and #8.2 displayed lower carbon concentrations in their stems compared to wild type (Fig. 12A,B). This observation prompted us to focus further on stem development in *OEP24.1* CRISPR mutants and overexpressing lines.

**Figure 12.**
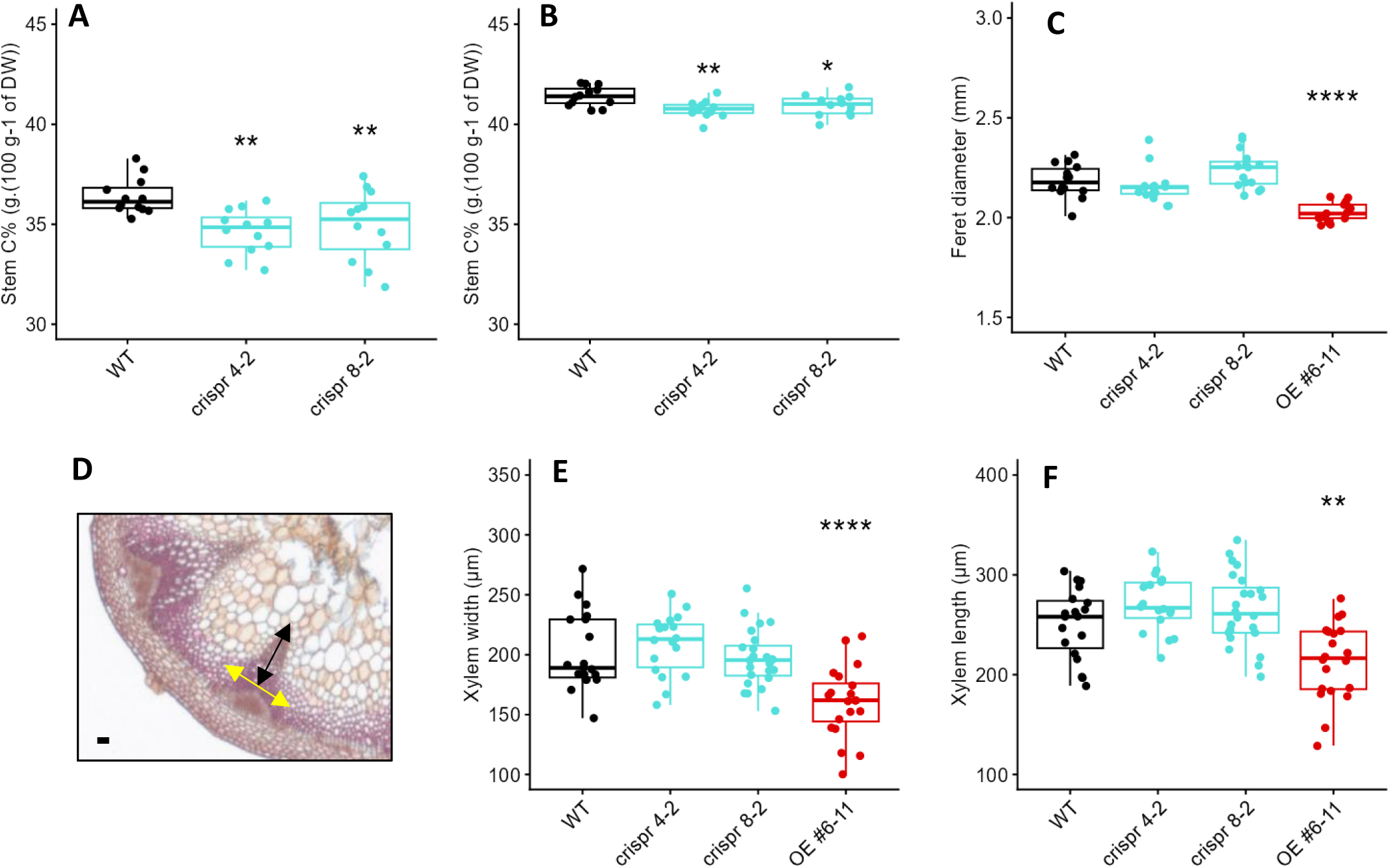
Stem carbon concentration is altered in OEP24.1 CRISPR lines while stem morphology, and xylem anatomy are altered in OEP24.1 over-expressing lines. (A,B) Stem carbon concentrations (C%) measured at harvest on wild-type (WT) and CRISPR mutants (4-2 and 8-2) grown under high-nitrate (A) or low-nitrate (B) conditions (n=12; T-tests; ** P<0.01; Supplemental Fig. S14. (C) Feret diameter measured on transverse stem section; n=15 to 18; T-test; **P<0.01, **** P<0.0001). (D) Representative transverse stem section stained using Alcian Blue/Safranin (2:1). Yellow and Dark arrows indicate xylem length and width respectively. Scale bar = 50 μm. (E) Xylem width and (F) Xylem length measured from transverse sections of wild type (WT), CRISPR mutants (4-2 and 8-2), and OE #6-11 over-expressing line as shown in (D), (n=15 to 18; T-test; **P<0.01, **** P<0.0001).

Changes in carbon concentration are known to impact cell wall composition, and there is evidence that cell wall composition affects stem diameter ^37^. Plants were grown in controlled growth chambers under long-day conditions until flowering to measure the diameter at the base of the primary stem. Stems were examined at 7 weeks after sowing, when flowers on the main stem were fully developed. While no obvious visual phenotype distinguished the #4.2 and #8.2 CRISPR mutants from the wild-type plants, OEP24.1 over-expressing line #6-11 displayed significantly thinner stems compared to the wild-type (Fig. 12C). Since a link between stem diameter and xylem development has also been previously established ^37^, we measured the radial and tangential lengths of xylem pole on stem transverse sections (Fig.12D-F), of the two CRISPR mutants and the #6-11 over-expressing line. Xylem radial and tangential lengths are used as indicators of cell proliferation and expansion, respectively. Interestingly, the xylem radial and tangential lengths were significantly smaller in the stems of the #6-11 over-expressing line, but were not significantly different from wild type in the mutant lines (Fig. 12E,F). This suggested that the overexpression of OEP24.1 affects both the cell proliferation and expansion in the xylem tissue. Stem diameter was then measured on the three independent OEP24.1 over-expressing line (#1-5, #6-11 and #9-6) using calypter, which confirmed that over-expressing lines have thinner stems (Fig. 13A). Cell wall composition was investigated using Fourier-transform infrared spectroscopy (FTIR). Mean spectra of the three OEP24.1 over-expressing lines show fingerprint regions that differ from WT (Fig. 13B) and t-values plotted against each wavenumber of the spectrum indicate significant changes (Fig. 13C), for the wavenumbers corresponding to the lignin and the cellulosic and hemicellulosic polysaccharides (864 cm⁻¹, 941 cm⁻¹, 1130 cm⁻¹, 1370 cm⁻¹, 1406 cm⁻¹, 1740 cm⁻¹; ^38–40^. More precisely, at 1740 cm^-1^, band specific to acetylated xylan ^41^ and at 1369 cm^-1^, corresponding to the deformation of C-H linkages in the methyl group *O*-acetyl moieties ^42^ suggest modifications in xylan acetylation. In addition, the hemicellulose peak at 1740 cm-1 was significantly smaller in all the overexpressing lines, suggesting less acetylated xylan (Fig. 13F). No change in the cellulose peak height (1050 cm^-1^, C-O vibration band) has been measured in #1-5, #6-11 overexpressing lines while a significant increase was measured in #9-6 (Fig. 13E). In addition, absorbance bands associated with G-type lignins (1140, 1250 and 1510 cm⁻¹; ^43^) exhibited significantly higher intensity in all the over-expressing lines than in the wild-type (Fig.13C). Nonetheless, the lignin height peak ratio (1510/1595 cm^-1^) which refers to the ratio between G-type specific band at 1510 cm^-1^ (C=C stretching) and G-type and S-type stretching vibration at 1595 cm^-1^ (C=C stretching of G-type and S-type and C=O), was significantly smaller in the OEP24.1 over-expressing lines compared to WT (Fig 13G). This suggests that even if G-type lignins are more abundant, their proportion compared to the S-type lignins are reduced. Altogether, these results show that the overexpression of OEP24.1 induces profound changes in the xylem secondary cell wall composition which could affect the cell wall rigidity and extensibility (cell wall richer in G-type lignins and less hemicelluloses) as well as the cell wall porosity since the reduction of hemicelluloses could affect their interaction with cellulose. Finally, consistent shifts were also observed in the NH₂-associated bands (approximately 1550–1650 cm⁻¹ ^44,45^), possibly reflecting changes in protein or amine-containing cell wall components. Overall, these results show that the overexpression of OEP24.1 affects the composition of xylem secondary cell walls.

**Figure 13:**
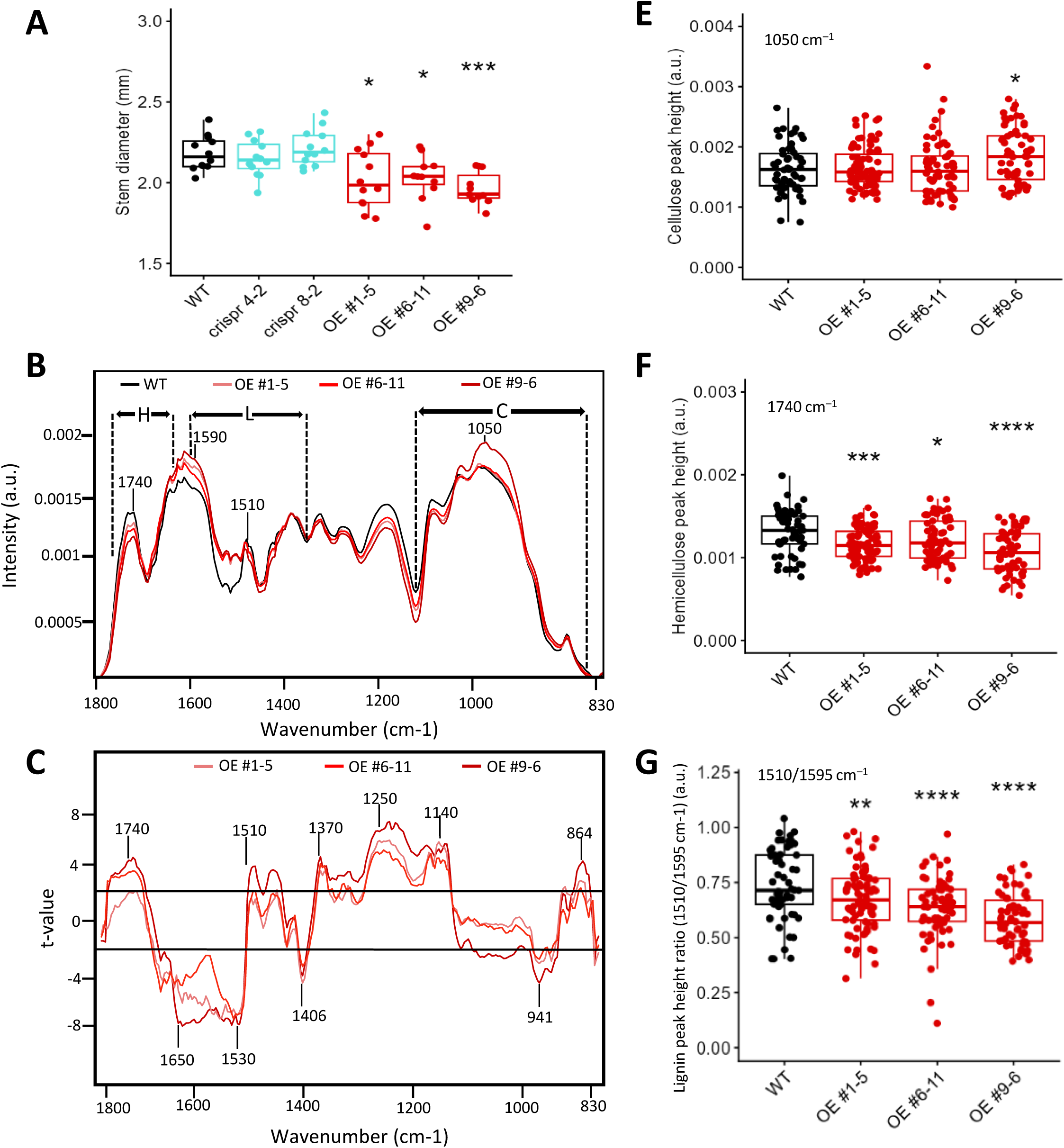
The composition of the xylem secondary cell wall is altered in OEP24.1 over-expressing lines. (**A**) Stem diameter was measured at the base of the primary stem using caliper on wild-type (WT), CRISPR mutants (4-2 and 8-2), and the three OEP24.1 over-expressing lines (OE #1-5; #6-11; #9-6) grown under long days and high nitrate conditions (n=12; T-test; *P<0.05, ** P<0.01, *** P<0.001). FTIR spectra were acquired on the xylem tissues from the transversal sections of the basal inflorescence stem. All spectra were baseline-corrected and area-normalized in the range 1800–800 cm–1. (B) Average FTIR spectra were generated from 60, 64, 83, and 62 spectra for the wild-type, OE#1-5, OE#6-11, and OE#9-6 plants (n=3 plants), H=Hemicellulose, L= Lignin and C=Cellulose. (C) Comparison of FTIR spectra obtained from xylem cells of the OE#1-5, OE#6-11, and OE#9-6 overexpressing lines. A Student’s t test was performed to compare the absorbances for the wild-type (WT) and the three over-expressing lines. The results were plotted against the corresponding wavenumbers. t-Values (vertical axis) between –2 and + 2 correspond to nonsignificant differences (P<0.05) between the genotypes tested (n = 3). t-Values above + 2 or below –2 correspond to, respectively, significantly weaker or stronger absorbances in the mutant spectra relative to WT. (E) Boxplots of the cellulose peak height (C–O vibration band at 1050 cm –1). (F) Boxplots of hemicellulose peak height (C–O and C–C bond stretching at 1740 cm–1). (G) Boxplots of lignin peak height ratio (1510/1595 cm–1). The lines represent median values, the tops and bottoms of the boxes represent the first and third quartiles, respectively, and the ends of the whiskers represent maximum and minimum data points. The boxplots represent values (shown as colored dots) from 60, 64, 83, and 62 spectra for the wild-type, OE#1-5, OE#6-11, and OE#9-6 plants (n=3 plants; T-test; *P<0.05, ** P<0.01, *** P<0.001, **** P<0.0001).

## DISCUSSION

### Piecemeal autophagy of chloroplasts

The turnover of chloroplast proteins is essential for maintaining the function of this organelle. From biogenesis to decay, chloroplast quality control is primarily governed by prokaryotic-type proteases that degrade internal chloroplast proteins. The turnover of outer envelope proteins, such as the TOC proteins involved in the protein import translocase, is mediated by the CHLORAD machinery, which involves the ubiquitination of outer envelope proteins and their subsequent degradation in the cytosol by the 26S proteasome ^46^. The E3 ubiquitin ligases SP1 and SP2 (suppressors of ppi1) are key components of this process. Additionally, the PUB4 (Plant U-box4) E3 ubiquitin ligase has been shown to facilitate the quality control and degradation of chloroplast proteins under dark/light stress conditions ^47^.

Mechanisms involved in chloroplast decay and dismantling during developmental senescence likely differ from ubiquitin/proteasome-mediated chloroplast protein quality control and operate independently. Wada et al. (2009) demonstrated that the transfer of Rubisco-containing bodies (RBCs), whole chloroplasts, and stroma-targeted DsRed markers to the vacuolar lumen is autophagy-dependent during dark-induced leaf senescence. The induction of autophagy by dark stress and its contribution to Rubisco degradation were further confirmed by Ono et al. (2013). Colocalization of stroma-targeted DsRed and GFP-ATG8 in chloroplast protrusions and in aggregates within the vacuolar lumen underscores the role of macroautophagy in the piecemeal degradation of chloroplasts ^17,18^. The roles of microautophagy and macroautophagy in the degradation of entire UV-B-damaged chloroplasts were elucidated by Izumi et al. (2017) and Nakamura et al. (2018) ^12,32^. Using live-cell imaging of chloroplast morphology in leaves undergoing sugar starvation-induced senescence, Izumi et al. (2024) observed the formation of budding structures emerging from chloroplasts in wild-type mesophyll cells ^28^. These budding structures were absent in autophagy-deficient mesophyll cells, suggesting that autophagosome biogenesis is required for chloroplast budding. In wild-type cells, buds were released from chloroplasts and incorporated into the vacuolar lumen as autophagic cargo. More recently, Kikuchi et al. (2020) ^48^ showed that neither *pub4* nor *sp1* mutations are involved in autophagy-dependent piecemeal chloroplast degradation. The *atg5.pub4* and *atg7.pub4* double mutants exhibit additive and more severe phenotypes compared to single mutants, including increased ROS production and accelerated leaf senescence. Detection of ubiquitinated proteins indicated that both PUB4-associated ubiquitination and autophagy synergistically contribute to chloroplast protein degradation in senescing Arabidopsis leaves.

### Autophagy receptors for piecemeal chloroplast degradation

Organelle-selective autophagy is thought to be controlled by receptor proteins that recognize target organelles and act as a bridge between the organelles and the isolation membrane, where ATG8 is anchored ^49^. The presence of receptors at the chloroplast outer membrane capable of interacting with NBR1 or ATG8 was hypothesized, and several ATG8-interacting candidates were predicted using in silico tools ^50^. However, none of these candidates have been shown to play a role in RBC targeting for piecemeal chloroplast degradation or in chlorophagy. Nevertheless, NBR1 has been found to associate with photodamaged chloroplast membranes, coating both the surface and interior of chloroplasts. This coating is followed by the engulfment of damaged chloroplasts into the vacuolar lumen via a process independent of the macroautophagy machinery, likely related to microautophagy ^51^. The authors demonstrated that the delivery of NBR1-decorated chloroplasts into vacuoles depends on the ubiquitin-binding UBA2 domain of NBR1 but is independent of the ubiquitin E3 ligases SP1 and PUB4. Here, we provide evidence that OEP24.1 functions as an autophagy receptor for piecemeal chloroplast degradation.

### OEP24.1: A Receptor for Autophagy-Dependent Piecemeal Chloroplast Degradation

Proteomic analyses conducted in our team revealed that OEP24.1 accumulates at high levels in the leaves of autophagy mutants under stress, in contrast to most chloroplast proteins ^29^ (Fig. 1). This accumulation may result from the induction of OEP24.1 gene expression and translation in autophagy mutants, although this was not supported by previous transcriptomic analyses ^52^, or from defects in OEP24.1 autophagy-dependent degradation, as observed for other cargoes and autophagy-related proteins ^4^. By studying the interaction of OEP24.1 with ATG8, its degradation upon autophagy induction, and its colocalization with ATG8, we have accumulated several lines of evidence indicating that OEP24.1 acts as a receptor in autophagy-dependent piecemeal chloroplast degradation. Consistent with a role in autophagy, the OEP24.1 gene is co-expressed with many autophagy genes according to ATTEDII (https://atted.jp/). OEP24.1 is not degraded in *atg5* mutants or in wild-type plants treated with concanamycin A. OEP24.1 binds to several ATG8 isoforms in a UIM/UDS-dependent manner. However, as OEP24.1 is embedded in the lipid membrane of the chloroplast outer envelope, the accessibility of its UIM site for ATG8 interaction raises questions, and the mechanism by which the OEP24.1 UIM site is exposed for ATG8 interaction remains to be elucidated. Notably, despite its uniform localization in the outer membrane of all plastids, regardless of plant tissue, colocalization of OEP24.1-GFP with RFP-ATG8 is only observed in budding structures emerging from chloroplasts and containing stroma proteins but not chlorophyll, which closely resemble the RBC bodies described in previous studies ^17,18,27,28^. Once released from chloroplasts, OEP24.1-GFP/RFP-ATG8 complexes behave as mobile autophagosomes that reach the central vacuole, where they are observed as OEP24.1-GFP-containing autophagic bodies upon concanamycin A treatment. These observations show that, when present at the chloroplast envelope, OEP24.1 interacts with ATG8 only at budding sites where autophagosomes are formed. Imaging of *atg5* mesophyll cells transformed with the OEP24.1-GFP construct revealed an overabundance of long stromules and the absence of OEP24.1 bodies in the cytosol and vacuolar lumen of *atg5* cells, demonstrating the dependence of OEP24.1 cellular trafficking on autophagy. Protein modifications that control the interaction of cargoes, adaptors, or receptors with ATG8 have been described in the literature. For example, the GSNOR enzyme requires nitrosylation to enable ATG8 binding ^53^, and DSK2 binding to ATG8 is conditional upon phosphorylation by BIN2 ^54^. Whether OEP24.1 is modified or processed at stromule budding sites, and whether this facilitates its interaction with ATG8, remains to be explored.

### Function of OEP24.1 in chloroplast and metabolism

OEP24.1 is a pore-forming protein initially identified in Pisum sativum (pea) as PsOEP24 ^55^. This β-barrel protein is ubiquitously expressed in plants but is most abundant in Arabidopsis seeds during maturation. Arabidopsis also encodes a closely related homolog OEP24.2 (At1g45170), which exhibits an opposite expression pattern. Although the functions of OEP24.1 and OEP24.2 remain unclear, *in vitro* reconstitution of pea PsOEP24 in proteoliposomes showed that the channel it forms is slightly cation-selective but highly conductive ^56^. PsOEP24 was reported to form homodimers with a 2.5- to 3-nm-wide pore that could facilitate the transport of triose phosphates, hexose phosphates, sugars, ATP, phosphate (Pi), and dicarboxylates ^55^. Authors showed that OEP24 allows the passage of glycine, valine, arginine and glutamate, whereas transport of aspartate was markedly slower. Another study demonstrated that pea PsOEP24 can functionally replace mitochondrial VDAC in yeast. In Arabidopsis, it remains unknown whether heterodimers of OEP24.1 and OEP24.2 exist. Co-expression network analysis (ATTED II) indicates that OEP24.1 is co-expressed with genes involved in amino acid synthesis, although its potential role in amino acid diffusion has not been tested. In pea, PsOEP24 was found to be uniformly distributed in the plastids of all tissues. Similarly, in Arabidopsis, our confocal imaging revealed OEP24.1-GFP localization at the plastid envelope in roots, hypocotyls, epidermis, and in mesophyll cells.

Phenotyping of *oep24.1* CRISPR mutants and *OEP24.1* overexpressors did not reveal as strong phenotypes as observed in the mutants of the core autophagy machinery. Nevertheless, these lines exhibited phenotypes related to carbon concentrations and allocation at the whole plant level, which may be linked to a role of OEP24.1 in metabolite flux between chloroplast and cytosol ^55^, and/or to its role as autophagy receptor in chloroplast quality control ^57,58^. The relatively mild phenotypic effects observed here suggest that OEP24.1 is neither the sole chloroplast receptor nor the only channel mediating metabolite efflux from the chloroplast.

## MATERIAL AND METHODS

### Plant material

Arabidopsis thaliana ecotype Col-0 was used as wild-type for all experiments. The *atg5* mutant (atg5-3, SALK_020601), *sid2* (sid2-2; ^59^) and *sid2.atg5* double mutant have been previously characterised by Yoshimoto et al. (2009) ^60^. The OEP24.1 CRISPR mutants (#4.2 and #8.2), the OEP24.1 over-expressors (#1-5, #6-11, #9-6) and the plants carrying GFP-ATG8e and OEP24.1-GFP fusions were produced in this study (see below).

### Plant Growth conditions

For nitrogen remobilisation assays, plants were cultivated in a growth chamber with short day conditions (8 hours of light and 16 hours of darkness) for 55 days after sowing in order to promote rosette development. Then, plants were transferred in long day conditions (16 hours of light and 8 hours of darkness) to induce flowering. Hygrometry was maintained at 65%. Plants were cultivated under low nitrogen nutrition (2 mM nitrate) or under high nitrogen nutrition (10 mM nitrate) as described in Masclaux-Daubresse and Chardon (2011) ^61^.

For stem phenotyping, plants were cultivated in a growth chamber with long day conditions (16 hours of light and 8 hours of darkness) for 49 days after sowing, stem was 40cm long. Hygrometry was maintained at 65%. Plants were cultivated under high nitrogen nutrition (10 mM nitrate) condition.

For confocal microscopy, seedlings were cultivated in vitro on solid MS/2 +1% sucrose agar media for 10 days. Then seedlings were transferred to (i) +NC liquid media (MS/2 + 1% sucrose) or (ii) -NC starvation liquid media (3 mM MES-NaOH; pH 5.5), or (iii) - NC liquid media with 1 μm concanamycin A, (CA; inhibitor of vacuolar H+-ATPase activity and proteolytic degradation).

### Plasmid constructs and plant transformation

All the plasmids used in this study are listed in supplemental Table S1. Plant transformations were performed via *Agrobacterium tumefaciens* (GV3101::pMP90) using the floral dipping. To create the OEP24.1 CDS without stop, the OEP24.1 CDS was amplified from the U15871 commercial clone, by a first PCR using the gtwOEP24 Forward and Reverse primers (Table S2). The product of this first PCR was used as a matrix for a second PCR amplification that used the U3 and U5 primers (TableS2). The product of the second PCR was cloned into the pDONOR207 vector via a BP Gateway reaction. The OEP24.1 CDS was then transferred from U15871 into the pMDC32 and pUBN-nYFP-Dest vectors and the OEP24.1 CDS without stops was transferred from the pDONR207 into the pUBC-GFF Dest vectors using LR Gateway reactions to obtain the p35S::OEP24.1, the pUbi::nYFP-OEP24.1 and the pUbi::OEP24.1-GFP vectors. Similarly, via an LR reaction, the ATG8e CDS was transferred from pDONOR207 (^62^) into the pH7WGR2 vector to obtain the p35S-RFP::ATG8e and ATG8d CDS was transferred from G22544 (commercial clone from ABRC) into pUBN-cYFP-Dest vector to obtain the vector pUbi:cYFP-ATG8d. All resulting constructs (see Table S1) were used for plant expression, subcellular localization, and BiFC experiments.

### Plasmid constructs for Y2H

For Y2H, the OEP24.1 CDS was transferred from the pDONOR207 to the pGAD vector and all the ATG8 CDS (Table S1) were subcloned into the pGBK vector using the LR gateway reaction. ATG8e and ATG8d sequences carrying mutations in the LDS site were synthetized by GeneArt Life Technologies (Thermo Fisher Scientific, Germany). To obtain ΔLDS-ATG8e and ΔLDS-ATG8d mutated isoforms, the Y_51_L_52_ residues were substituted with two alanines (TACTTG (ATG8D), TACCTT (ATG8E)-> GCTGCT) in the pDONR221 vector and subsequently subcloned into the pGBKT7 vector. Since both pDONR221 and pGBKT7 share the same antibiotic resistance cassette, a plasmid digestion was performed to remove the kanamycin resistance marker. Briefly, pDONR221 constructs were incubated overnight at 37 °C with the restriction enzymes *PvuI* and *ApaLI* in Tango buffer, generating a fragment containing the ATG8E sequence flanked by attL1/attL2 sites but lacking the kanamycin resistance cassette (Table S1). The digested fragment was separated by agarose gel electrophoresis, purified using the Gel and PCR Clean-Up Kit, and introduced into pGBKT7 by Gateway recombination. ATG8e and ATG8d sequences carrying mutations in the UDS site were generated by site-directed mutagenesis using the QuickChange PCR method. The I_78_F_79_I_80_ residues in ATG8e and ATG8d were substituted with 3 alanines (ATCTTCATT-> GCTGCTGCT) and subsequently cloned into the pGBKT7 vector via Gateway recombination. Each mutation was introduced with a single primer pair (Table S1). All the constructs in the vectors listed in supplemental Table S1 were verified by sequencing.

### Generation and selection of CRISPR mutants and OEP24.1 overexpressors

The guide RNA targeting the OEP24.1 sequence (ACTGACAAAACCAGTGGAAT and GTCGCCGGTCCTACTTTGAC) were designed using the CRISPOR software (^63^). The vector for plant transformation containing the guide RNAs and the CRISPR-Cas9 was synthetized by Eurofins Genomics using the pRMCas 9 plasmid. After transformation of Col-0 with this vector, ten homozygous lines of several primary transformants were selected on the basis of their hygromycin resistance and single insertion segregation rate. Sequencing using the crisprOEP-Forward and crisprOEP-Reverse primers (Table S2) was performed to identify mutant lines. For further study, we selected the OEP24.1 #4.2 and #8.2 mutated lines presenting deletion and insertion/deletion respectively, between the 2 sequence targets. These modifications of the OEP24.1 sequence in the #4.2 and #8.2 mutants result in the translation of truncated proteins of less than 20 amino acids (Supplemental Figure S1). The p35S::OEP24.1 vector was used to transform Arabidopsis and obtain the three homozygous over-expressor lines (#1-5, #6-11, #9-6) selected on the basis of their hygromycin resistance and single insertion segregation rate. After transformation of Col-0 with the pUbi::OEP24.1-GFP vector, several homozygous lines (#1-4, #2-11, #4-9, #5-1, #8-8, #12-5) were selected on the basis of their basta resistance and single insertion segregation rate. The line the pUbi::OEP24.1-GFP #12-5, providing good signals in confocal microscopy, was chose to be transformed by p35S-RFP::ATG8e vector. After transformation two homozygous lines (#3G, #10C) were selected on the basis of their hygromycin resistance and single insertion segregation rate.

### RT-qPCR

Total RNA and RT-qPCR were performed according to Di Berardino et al. (2018) ^64^ in order to verify the higher expression of *OEP24.1* in overexpressing lines. All the primer used are listed in Table S2. EF-1α (At5g60390) and ACT2 (AT3G18780) were used as reference genes.

### Confocal microscopy analyses

Confocal microscopy was used for both *in situ* localisation in Arabidopsis or *Nicotiana benthamiana* after agroinfiltration, or for BiFC in agroinfiltrated *N. benthamiana*. Fluorescence signals from chlorophyll (auto-fluorescence) and the eGFP, mRFP, and eYFP reporters were imaged using a confocal scanning microscope (Leica TCS-SP5). The manufacturer’s instructions. Line sequential scanning mode with dual-channel observation was applied to avoid possible bleed-through of signals from two fluorophores and chlorophyll. All images were processed by using ImageJ software (https://imagej.net/Fiji). We then chose a representative optical section for presentation of the data in the manuscript. An ImageJ/FIJI plugin to rapidly build scientific figures, EZFig, was used ^65^.

### Agrobacterium tumefaciens-mediated transient expression in N. benthamiana leaves

Agrobacterium strain C58C1pMP90 was transformed by electroporation with the binary vectors constructed to observe *in situ* localisation or for BiFC (see supplemental Table S1). Agrobacterium strain containing the P19 plasmid were also used. After overnight culture at 28°C in LB medium containing antibiotics (50 µg/mL rifampicin, 50 µg/mL gentamycin, and 50 µg/mL spectinomycin), agrobacteria were centrifuged and resuspended in infiltration buffer (10 mM MgCl2 and 10 mM MES pH 5.7) at 0.1 OD600 for each construct. Agrobacteria infiltration was performed at the abaxial face of leaves from 4-week-old plants using a syringe, 2 to 3 d before imaging.

### In planta co-immunoprecipitation (CoIP) analysis

CoIP and IP assays were performed as described by manufacturer (ChromoTek) with minor modifications. For Co-IP, ten-day-old plants carrying the p35S:RFP-ATG8e and pUbi::OEP24.1-GFP seedlings were subjected or not to dark stress. For dark stress, seedlings were placed in complete darkness for 24 h prior to harvest. For heat stress, seedlings were first exposed to a heat shock at 37 °C for 1.5 h, followed by a recovery phase at 23 °C for 1.5 h, and then subjected to a second heat shock at 44 °C for 45 min and harversted 48 h later. Seeding material were collected in liquid nitrogen, grinded and total proteins were extracted using an extraction buffer containing 10× RIPA lysis buffer (Millipore, https://www.merckmillipore.com/FR/fr), 100 mM PMSF, 1.5% DDM, and a 25× protease inhibitor cocktail (Roche; https://www.roche.fr/topic/science). Total proteins were incubated with anti-GFP antibody decorated microbeads (ChromoTek GFP-Trap™, Thermo Fisher Scienfic, Illkirch, France) or with anti-RFP antibody decorated microbeads (ChromoTek RFP-Trap™, Thermo Fisher Scienfic, Illkirch, France) overnight at 4°C on a rotary shaker. Washing steps were performed by following the manufacturer’s instruction. The proteins were eluted using 1× Laemmli buffer to release the immunoprecipitated proteins.

### Immunoblot analysis

Total protein extraction, fractionation, and immunoblotting were performed as previously described ^66^. Briefly, proteins were separated by SDS-PAGE on 4–20% denaturing polyacrylamide gels (Mini-Protean TGX Stain-Free Gels; Bio-Rad, Marnes la Coquette, France) and electroblotted onto nitrocellulose membranes (Trans-Blot Turbo RTA Mini 0.2 µm Nitrocellulose Transfer Kit #1704270; Bio-Rad, Marnes la Coquette, France). Proteins of interest were detected using rabbit polyclonal antibodies produced by PhytoAb company (https://www.phytoab.com/) raised against the specific peptide *KASIKGKYDTDKTSG* (dz297) from OEP24.1; the rabbit polyclonal antibodies raised against GFP (ab290, Abcam, https://www.abcam.com/en-us), RFP (ABIN129578, Antibodies-Online, Aachen, Germany), and ATG8a (anti-APG8a, ab77003, Abcam).

Blots were imaged at different exposure times, sufficient to obtain distinct bands without pixel saturation. Band intensities were quantified using Image Lab 6.1 software by calculating the surface area under each band profile. The relative intensities of the free GFP band and the OEP24.1–GFP fusion protein were then used to calculate ratios indicative of autophagy-mediated degradation.

### Protein-Protein Interaction Studies in Yeast

Y2H assays were carried out using the Matchmaker GAL4 Two-Hybrid System 3 from Clontech as previoulsy described ^67^. For Y2H analyses, pGADT7- and pGBKT7-derived constructs were introduced into the *Saccharomyces cerevisiae* AH109 strain by lithium acetate transformation. Transformed yeast colonies were selected on synthetic dropout (SD) medium lacking leucine and tryptophan (SD–LW). Protein–protein interactions were assessed by growth on selective medium lacking leucine, tryptophan, and histidine (SD–LWH), supplemented with 3-amino-1,2,4-triazole (3-AT) at 1 mM or 5 mM of 3 independent transformed yeast colonies representative of 10 independent transformed yeast with the same plasmids.

### Stem Phenotyping

Plants were grown under long-day conditions (16 h light/8 h dark) for 49 days. For each plant, the height of the flowering stem was measured with a ruler, and the stem diameter was measured at the bottom part of the stem, using a digital caliper, before harvesting a 1- to 2-cm segment taken at the bottom part of the stem. The stem segments were embedded in 8% (w/v) agarose solution and sectioned with a VT100 S vibratome (Leica, https://www.leica-microsystems.com/). Inflorescence stem cross-sections were stained using FASGA staining solution ^37^ and were imaged under an Axio Zoom V16 microscope equipped with a Plan-Neofluar Z 2.3/0.5 FWD 10.6 objective (Zeiss, https://www.zeiss.fr/microscopie/). For each section, the diameter and xylem dimensions (length and width) of the inflorescence stem were measured using the FIJI software (https://fiji.sc/). The composition of the secondary cell wall of the xylem tissue was also determined on previous stem cross-sections by Fourier Transformed Infra-red spectroscopy using an FT-IR Nicolet iN (Thermo Fisher Scientific, https://www.thermofisher.com). Spectral acquisitions were done in reflexion mode on a 30 µm × 30 µm acquisition area targeting the xylem tissue as described in ^56^. Between 5 and 6 acquisition points sweeping the xylem tissue homogeneously were performed on 1 vascular bundle within a stem section, four individual stems section were analysed for one individual stem. Four individual inflorescence stems were analysed for each genotype. After sorting the spectra and correcting the baseline, the spectra were area-normalized and the different genotypes were compared to wild-type plants ^37^. The absorbance values (maximum height) of the major cellulose, lignin and hemicellulose bands in the fingerprint region (1,800–800 cm^−1^) were collected using the ‘HyperSpec’ library in R ^68^.

### 15N labelling and ^15^N, N and C measurements

To evaluate nitrogen remobilization, ^15^N, N and C concentrations and partitioning were measured and calculated according to Masclaux-Daubresse and Chardon (2011) ^61^. In brief, plants grown with sufficient or limiting nitrate supplies, were labelled with ^15^NO_3_^-^ for 24 h at vegetative stage and harvested after a long chase period when seeds were matured. Dry plants were then separated in rosette, stem and seeds to quantify 15N, N and C concentrations using elemental analyser and IRMS and calculate partitioning.

### In silico protein modelling and interaction prediction

Structural models were generated using AlphaFold v3 (DeepMind, 2024). The full-length amino acid sequence of the candidate protein OEP24.1 (TAIR: At5g42960; UniProt: Q9FJ57) was submitted to obtain its predicted fold. To explore its potential role as a porin, models were also generated in complex with over 50 different lipid ligands, which consistently supported its incorporation into a lipid bilayer environment. To assess possible interactions with autophagy-related proteins, we modeled the candidate protein OEP24.1 together with the nine Arabidopsis ATG8 isoforms ATG8a-i (Q8LEM4, Q9XEB5, Q8S927, Q9SL04, Q8S926, Q8VYK7, Q9LZZ9, Q8S925, Q9LRP7). For ATG8a-g, the C-terminal glycine-exposing cleavage by ATG4 was mimicked in silico by truncating the terminal residues downstream of the conserved glycine, thereby reproducing the processed form competent for PE conjugation. ATG8h and ATG8i, which are naturally processed, were used in their native sequence. Predicted interaction models were evaluated based on the predicted alignment error (PAE) and the interface pTM (ipTM) scores.

## Accession Numbers

The Arabidopsis Genome Initiative locus identifiers for the genes mentioned in this article are OEP24.1: At5g42960: ATG8 isoforms: At4g21980, At4g04620, At1g62040, At2g45170, At3g60640, At5g37640, At5g50230, At5g06420, At3g06420; ATG5: At5g42960

## Author contributions

CMD and AM designed the research and supervised LL PhD work. AM constructed the CRISPR mutants. JL constructed over-expression lines and protein fusions. LL performed plant phenotyping on mutants and over-expressing lines. RLH contributed to stem anatomy and cell wall composition. LL performed Y2H assays. BB and LL performed co-immunoprecipitations. AM and CMD performed confocal microscopy for co-localisations and BiFC. FC contributed to statistical analyses of phenotyping data. CMD wrote the article, with contributions from all authors. All authors approved the final version of the article.

## Supporting information

Supplementary Lambret et al.

## Acknowledgements

Authors thank Fabienne Soulay, Mégane Mahieu, Bastien Baudy and Luc Bachelet for technical help. Thanks to Amélie Bernard (LBM, CNRS, Bordeaux, France) for proof reading and discussions.

