## Supplementary Lambret et al. for "OEP24.1 involved in carbon allocation is a receptor of piecemeal plastid autophagy in Arabidopsis"

[£] authors contributed equally

* Corresponding authors

#### Short title: Role of OEP24.1 in autophagy and carbon allocation

The author(s) responsible for distribution of materials integral to the findings presented in this article in accordance with the policy described in the Instructions for Authors (https://academic.oup.com/plcell/pages/General-Instructions) is (are): Marmagne A. and Masclaux-Daubresse C. Authors (

### SUPPLEMENTARY MATERIAL

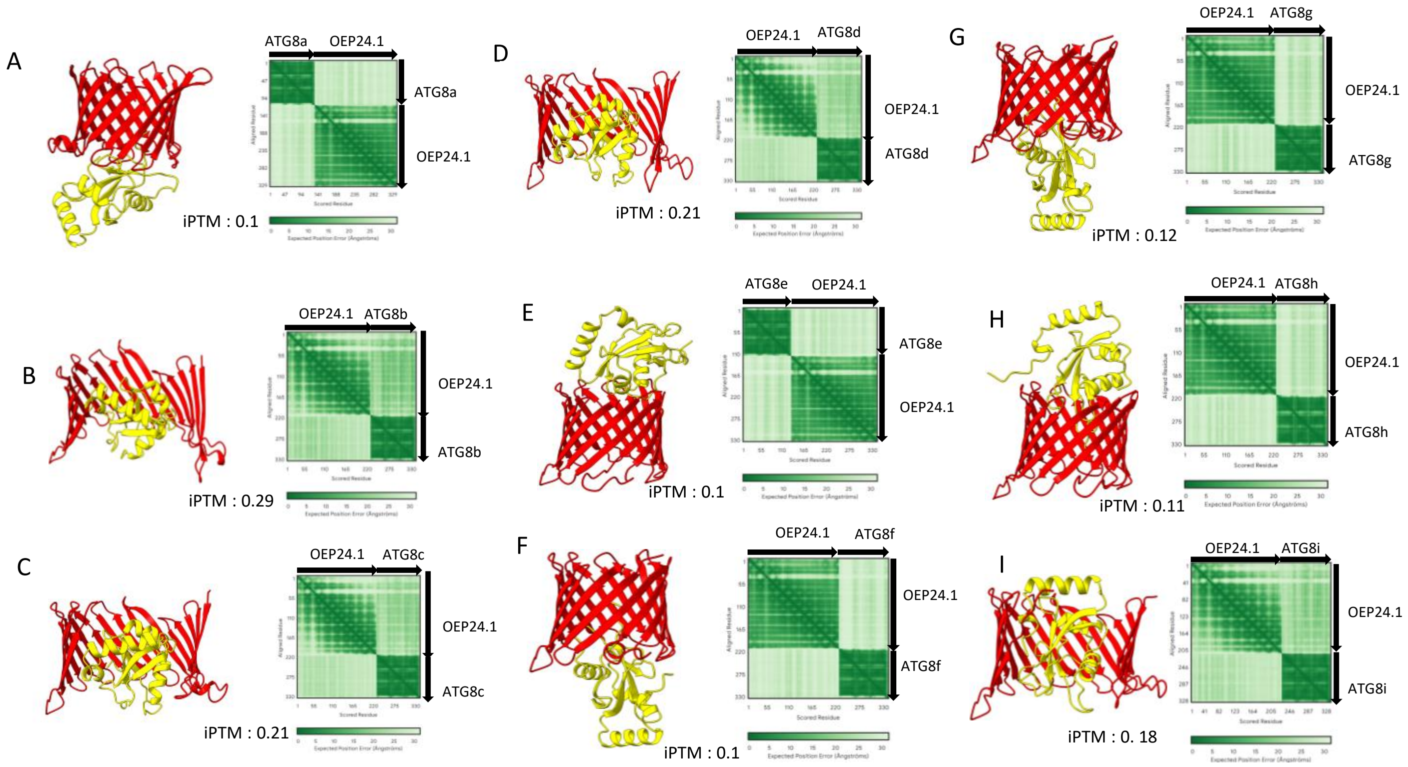

**Supplemental Fig.S1: Alpha Fold predictions for protein-protein interactions between OEP24.1 and the 9 ATG8a-i isoforms.** Predicted structural models of OEP24.1 (red) in complex with the nine ATG8 isoforms (yellow) were generated using AlphaFold3 (panels A-I corresponding to ATG8a-i isoform respectively). For each model, the corresponding Predicted Aligned Error (PAE) plots and the interface predicted TM-scores (iPTM) are shown.

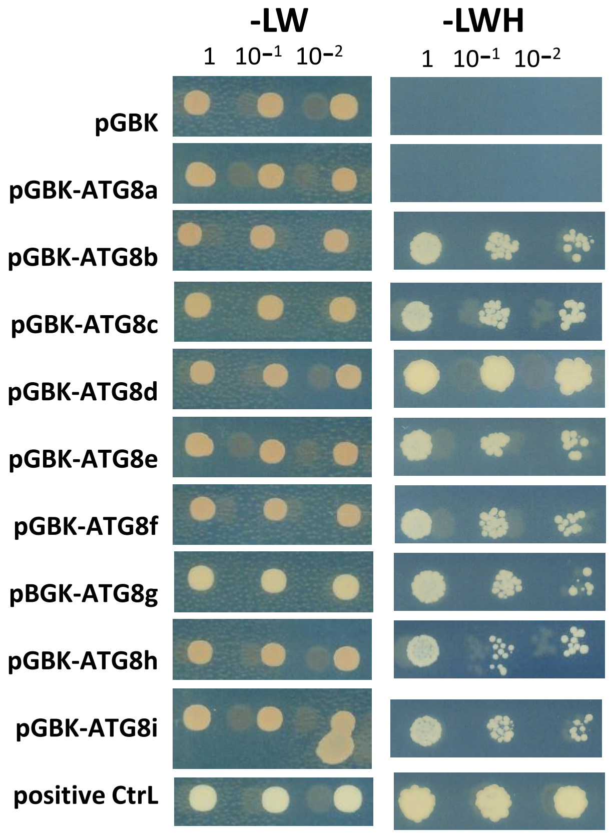

**Supplemental Fig.S2: OEP24.1 interacts with all the ATG8 isoforms except ATG8a**. ATG8a-i isoforms were used as bait (BD fusion) and OEP24.1 as a prey (AD fusion). Yeasts harboring both bait and prey expression vectors were grown on medium without His (-LWH) to check for interaction between both partners. Results show the growth of the 1,$10^{-1}$ and $10^{-2}$ dilutions of a representative clone after yeast conjugation. Experiment and yeast conjugation were repeated 3 times showing same results. Positive control consisted in ATG8g (BD) and NBR1 (AD) interaction. Negative control consisted in testing interaction between empty pGBK (BD) and OEP24.1 (AD).

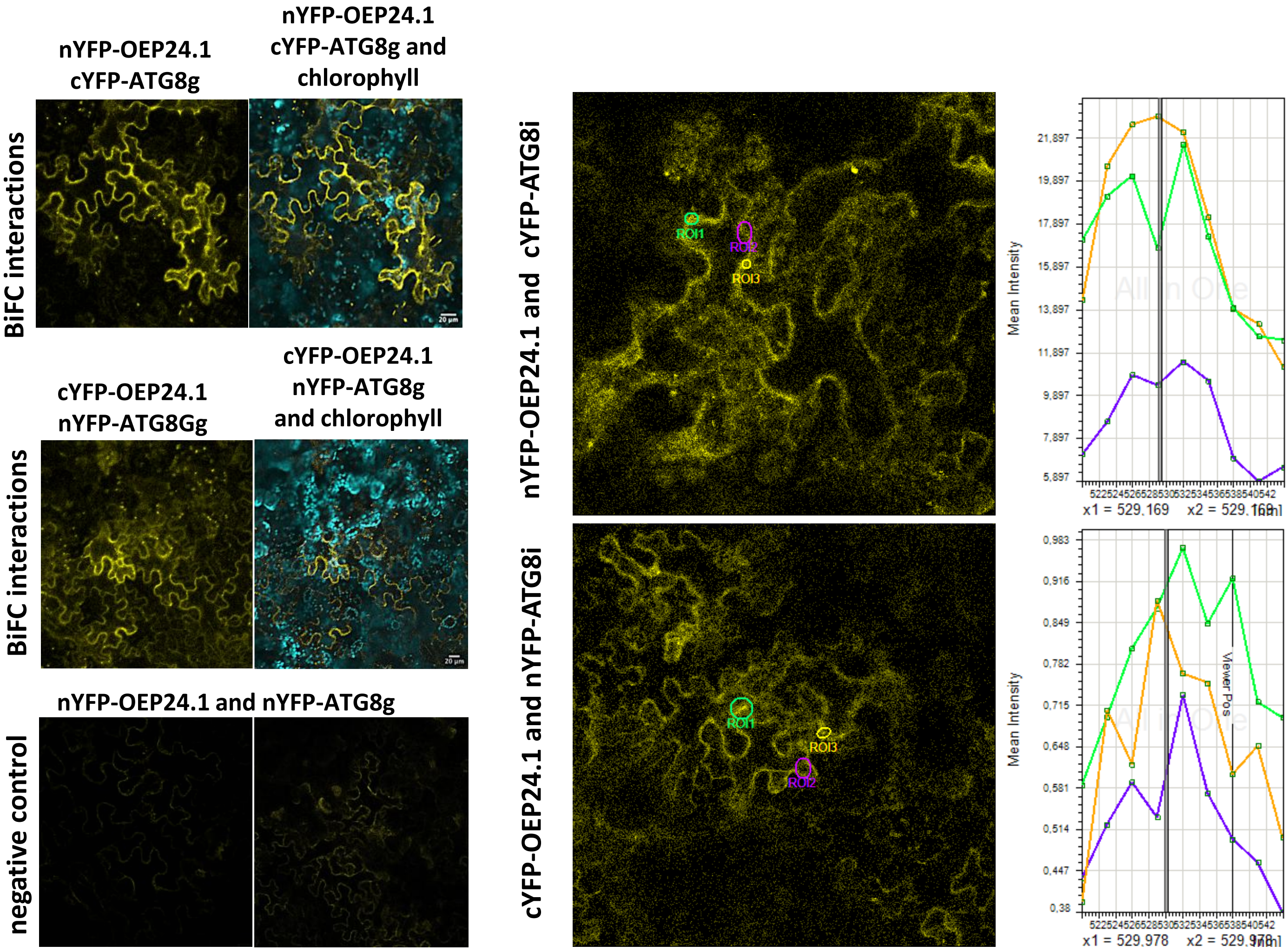

**Supplemental Fig. S3: OEP24.1 interacts with ATG8 *in planta*.**

BiFC was performed on leaves of mature *Nicotiana benthamiana* plants co-agroinfiltrated with the vectors pUBi::(n/c)YFP–OEP24.1, pUBi::(n/c)–ATG8i/g, and p35S::P19. Confocal imaging was performed by exciting the tissues with a 514 nm laser. Fluorescent signals were detected, and spectral analyses indicated emission peaks at 529 nm, characteristic of YFP fluorescence. The areas used to generate the emission spectra are indicated by circles, with their colours corresponding to the spectra shown on the right. Experiments were repeated three times (n = 3), yielding consistent results.

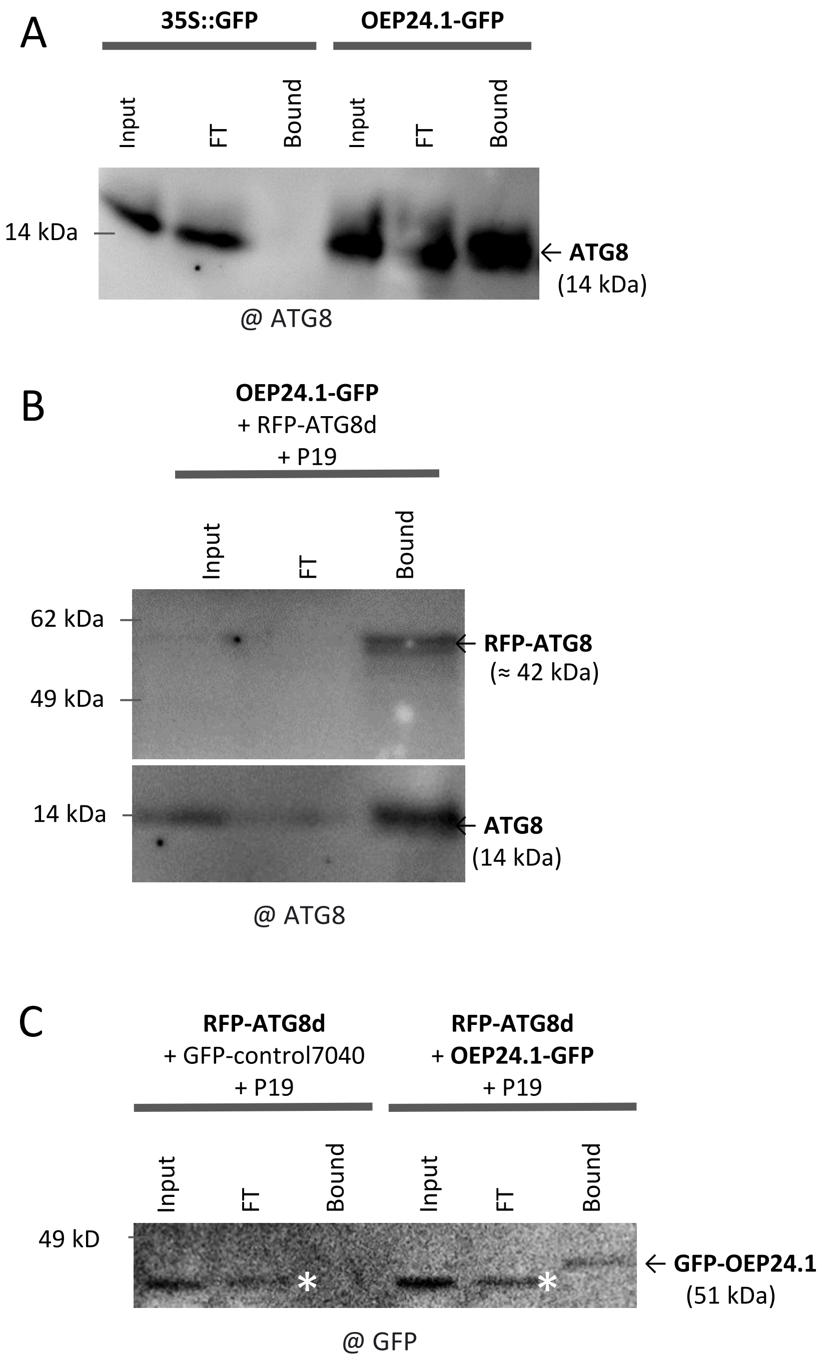

**Supplemental Fig.S4: Immunoprecipitation and co-immunoprecipitation in Arabidopsis and tobacco confirm interactions between OEP24.1 and ATG8.**

(A) Immuno-precipitations were performed by GFP-trap on Arabidopsis seedlings carrying p35S::GFP (control) or pUbi::OEP24.1-GFP constructs. Immunodetection of ATG8 was positive in the bound extract of OEP24.1-GFP plants and negative in the bound extract of the p35S::GFP control plants; (B) Co-IP was performed by GFP-trap on *Nicotiana benthamiana* leaves coinfiltrated with OEP24.1-GFP and RFP-ATG8d constructs. Both RFP-ATG8 and free ATG8 were detected in the bound extract. (C). Co-IP was performed by RFP-trap on *Nicotiana benthamiana* leaves coinfiltrated with RFP-ATG8d construct and OEP24.1-GFP construct (right) or GFP-7040 control (left; 7040 is used as a control protein known to be located in the chloroplast). Signal corresponding to the GFP-OEP24.1 molecular weight was detected in the bound extract odf the OEP24.1-GFP and RFP-ATG8d coninfiltrated tissues but not in the GFP-7040 and RFP-ATG8d coinfiltrated tissues. The white asterisk correspond to unspecific signal that is also present in this control.

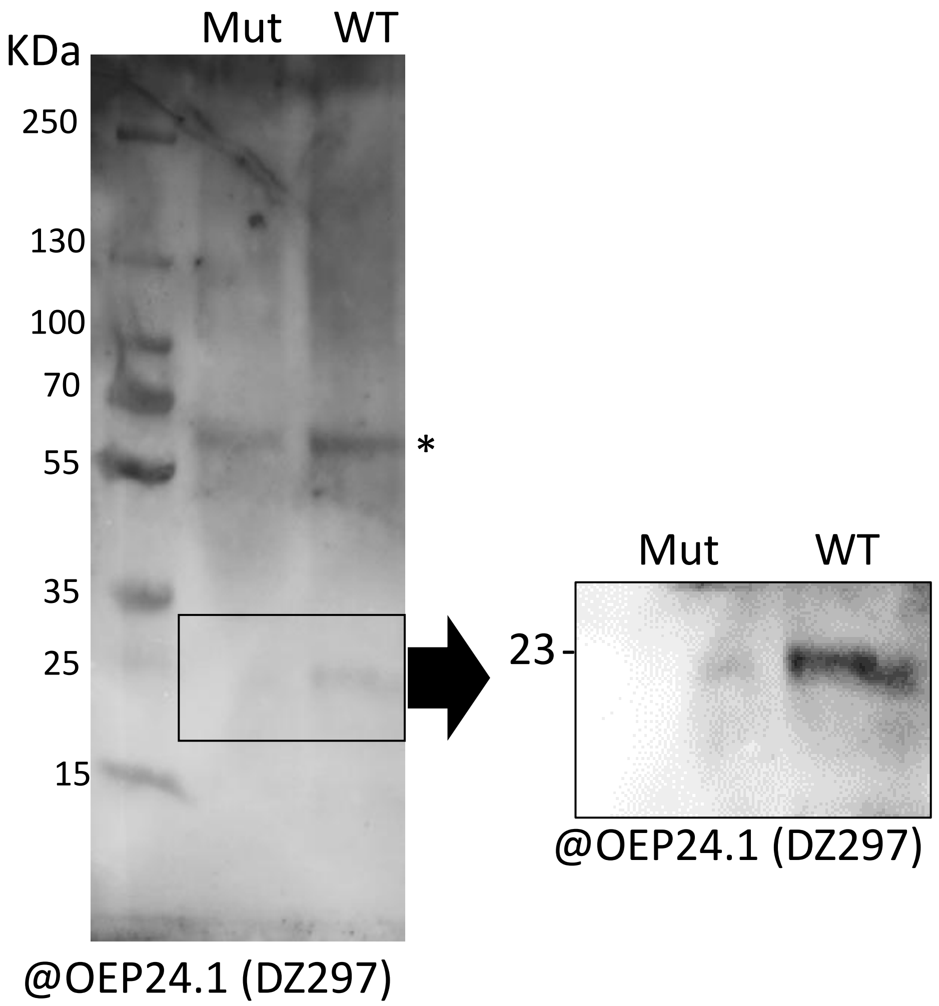

**Supplemental Fig.S5 : The DZ297 antibodies is specific of OEP24.1.** DZ297 antibodies were obtained from PHYTAB manufacturer (https://www.phytoab.com/) in rabbit after inoculation with the synthetic peptide KASIKGKYDTDKTSG. To verify specificity of these antibodies againts AtOEP24.1, equal amount of leaf soluble proteins (10 mg) were loaded in lane 1 (OEP24.1 4-3 mutant) and lane 2 (wild type) of acrylamide gel (4-20% acrylamide) for separation, transfered on membrane and blotted with DZ297. A band of a molecular mass of 23 KDa, which corresponds to OEP24.1 MW, was detected in WT extract only. Non specific signal is detectable between 55 and 70 KDa (*).

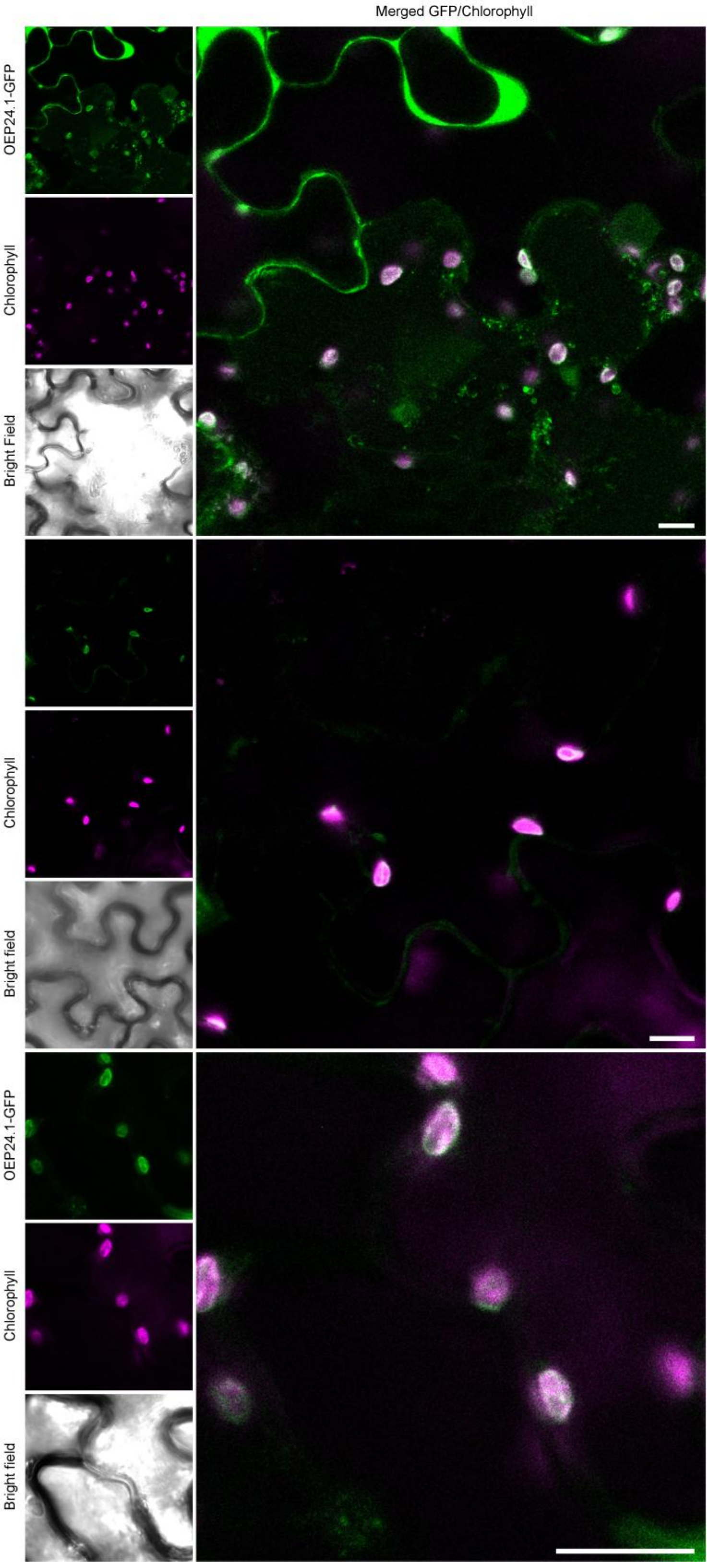

**Supplemental Fig.S6: OEP24.1 is present in at the chloroplast envelope in *Nicotiana benthamiana*.** *Nicotiana benthamiana* epidermis cells transiently expressing OEP24.1-GFP construct after agroinfiltration were analyzed by confocal microscopy. Three panels present different representative epidermis cells expressing the fusion, from different experiments (n=3). Green color shows the fluorescent signal of the OEP24.1-GFP protein fusion, magenta color shows chloroplast autofluorescence. The large image of each panel merges these two signals. Scale bars: 10 µm.

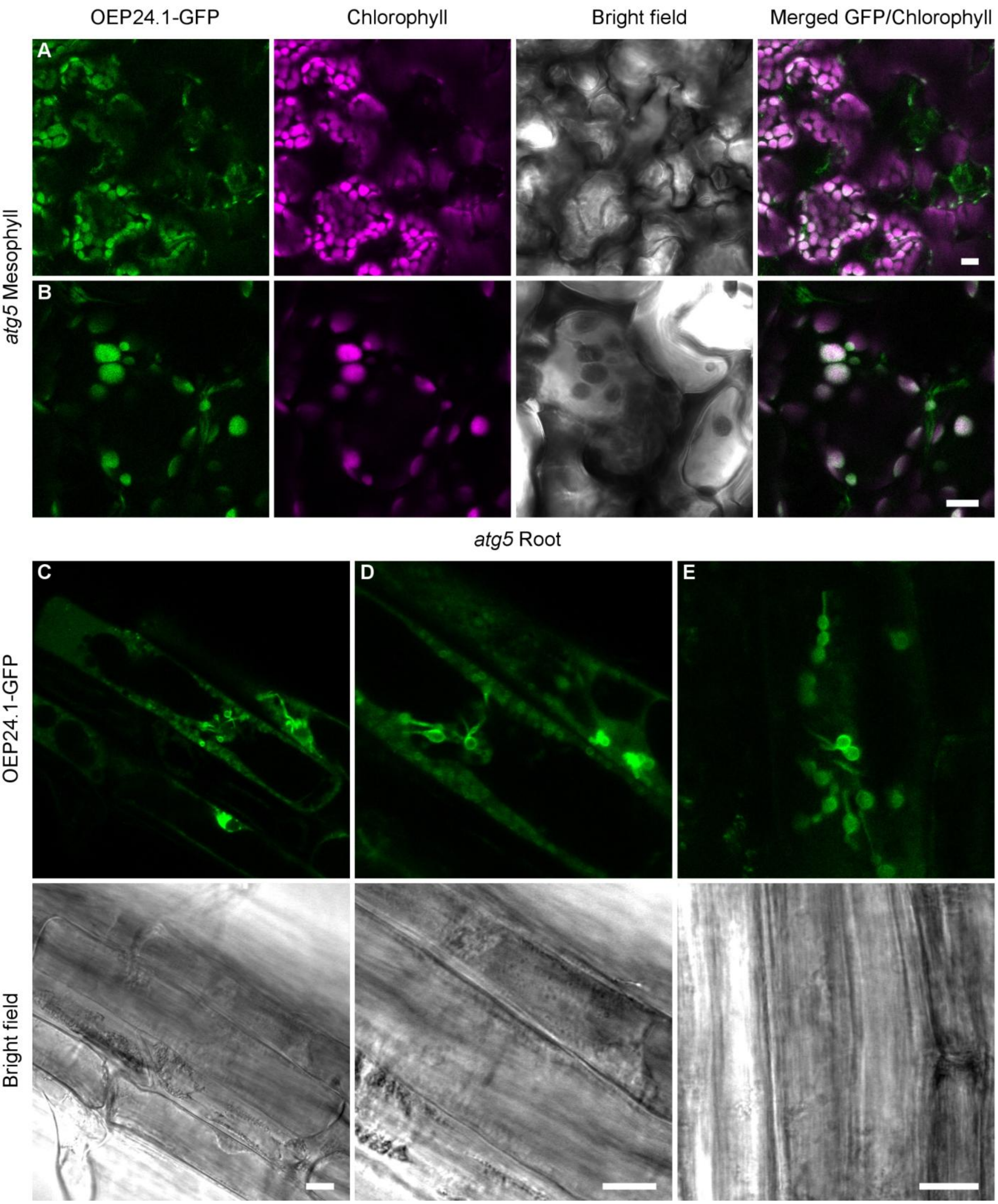

**Supplemental Fig. S7: OEP24.1 is localized at the chloroplast and etioplast envelopes in *atg5* mesophyll and root cells.** Leaves and roots of OEP24.1-GFP *atg5* seedlings were analyzed by confocal microscopy. (A,B) Like in wild type the fluorescence of OEP24.1-GFP was observed at the envelope of large and small chloroplasts in mesophyll et epidermis cells. (C-E) OEP24.1-GFP fluorescence was also observed in roots on small plastids exhibiting numerous stromules. Green color shows the fluorescent of the OEP24.1-GFP protein fusion, magenta color shows chloroplast autofluorescence. Bright field shows cell boundaries. Imaging was repeated (n>3). Scale bars: 10 mm.

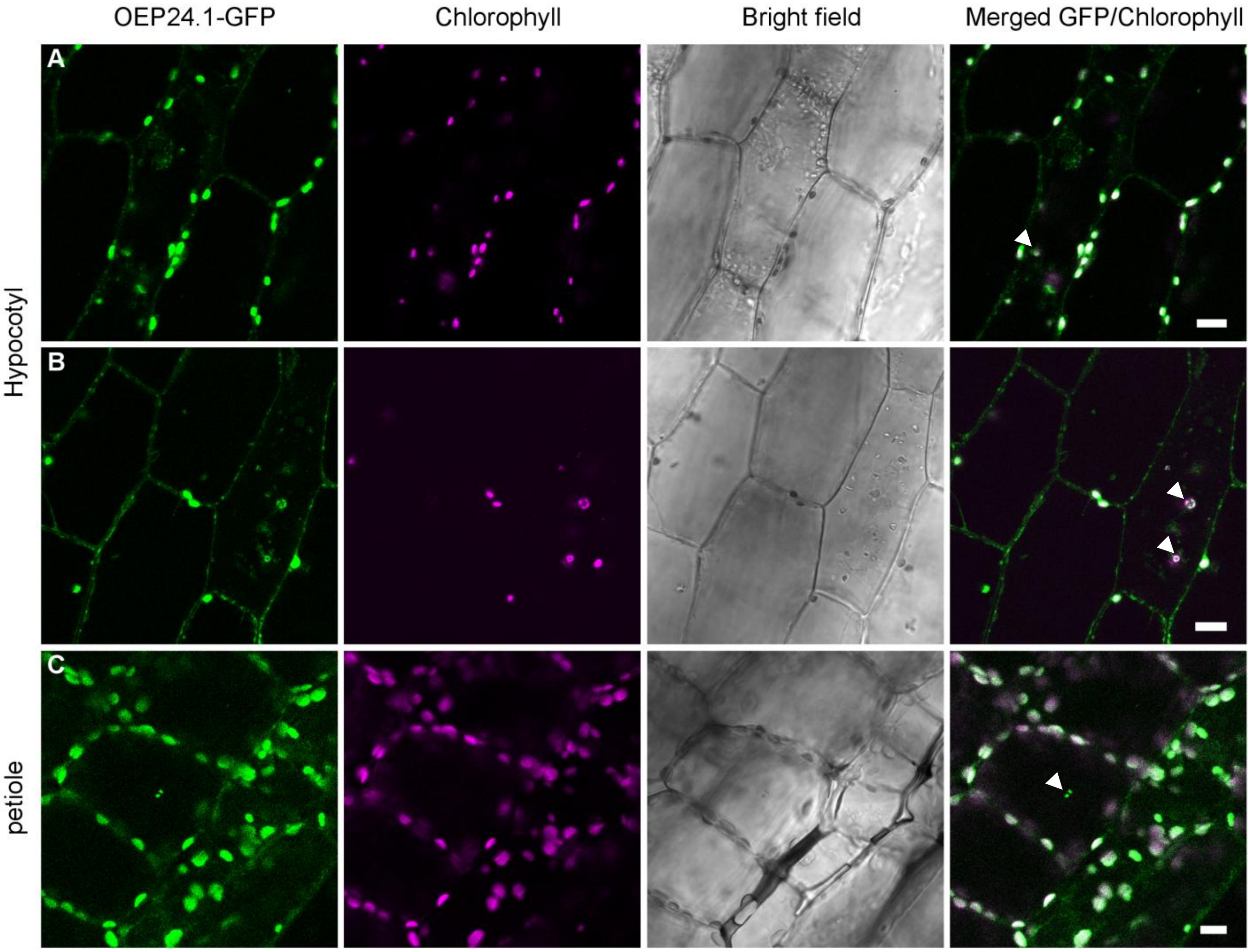

**Supplemental Fig.S8: Accumulation of OEP24.1 in aggregates and autophagic bodies in hypocotyl cells after concanamycin A treatment**. Plants carrying OEP24.1-GFP were treated with 1 µM concanamycin A, 24 hours before microscopic observation. GFP fluorescence signal was visualized in aggregates and autophagic bodies (white arrows; in the vacuole) the cells of hypocotyl and petiole by fluorescence confocal microscopy. scale bars: 10 mm.

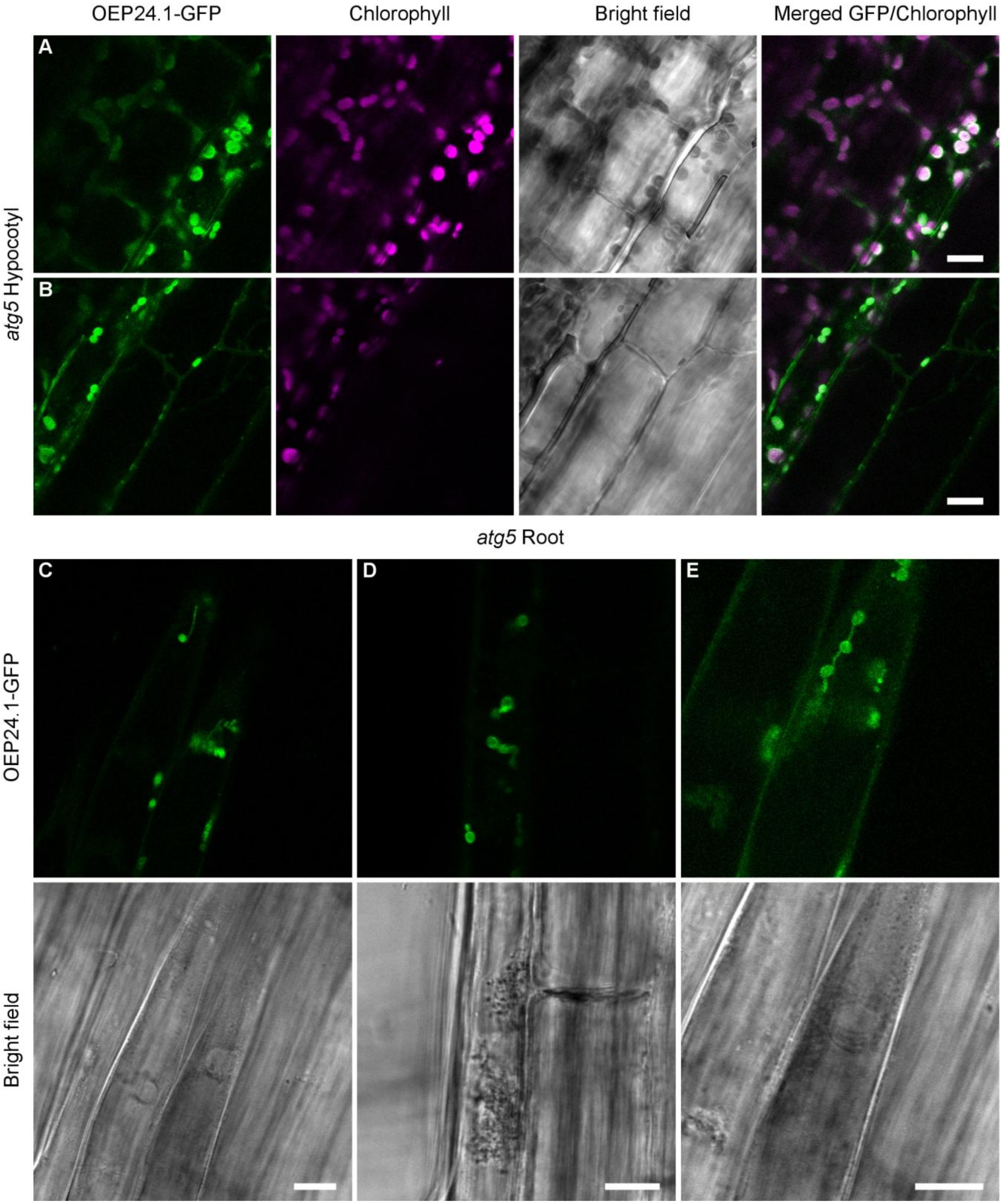

**Supplemental Fig.S9: Transfer of OEP24.1 to the vacuole upon concanamycin A treatment is autophagy dependent.** Leaves and roots of *atg5* seedlings expressing the OEP24.1::GFP were analyzed by confocal microscopy. Plants were treated with 1 mM concanamycin A 24 hours before microscopic observation. The GFP fluorescence was observed at the plastid envelopes but was absent from the vacuole lumen. (A-C): Representative images of hypocotyl cells. (C-E) Representative root cells. n=3. scale bars: 10 mm.

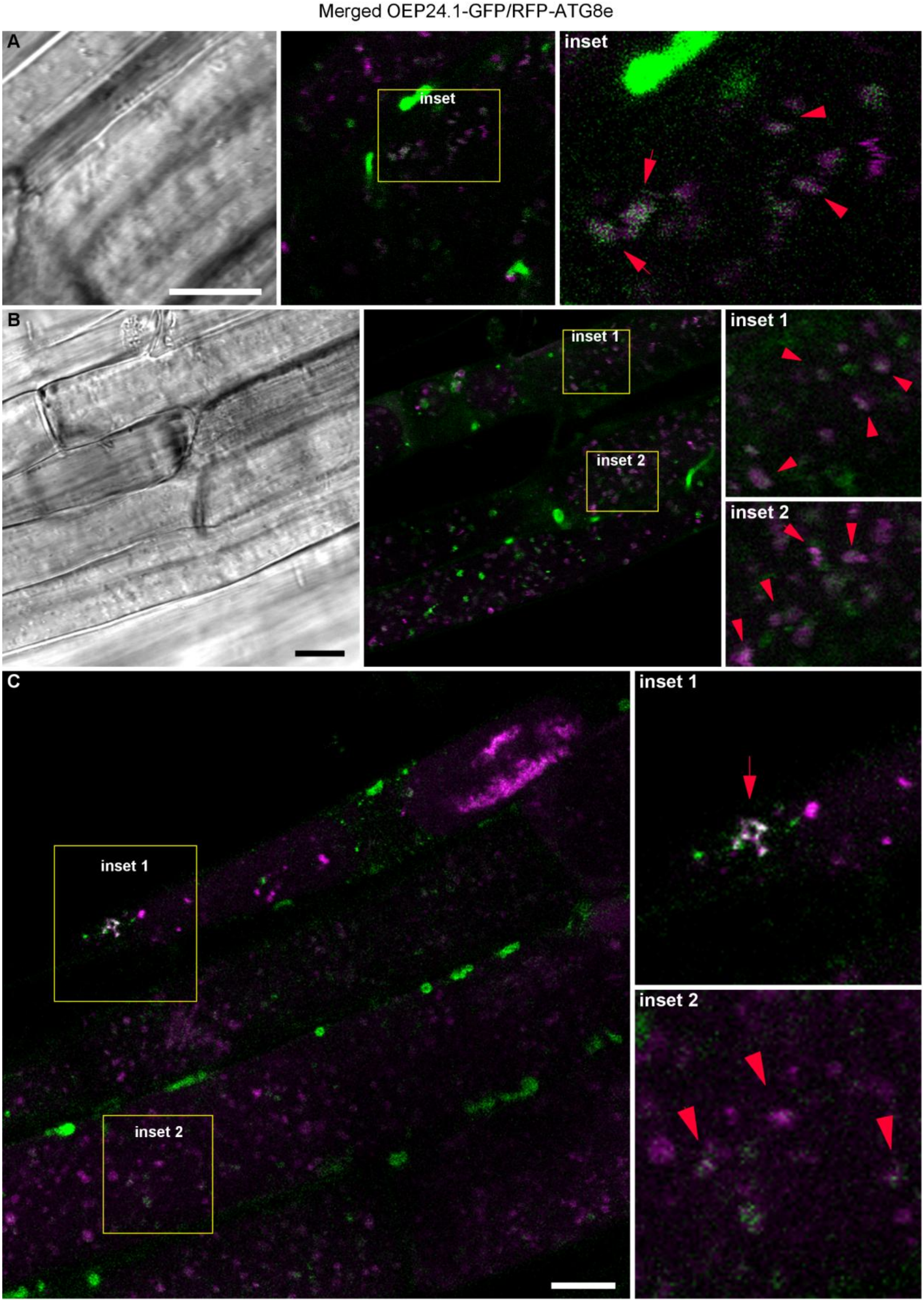

**Supplemental Fig.S10: OEP24.1 and ATG8e colocalize in aggregates and autophagic bodies of root cells after concanamycin A treatment**. Plants were treated with 1 µM concanamycin A, 24 hours before microscopic observation. GFP and RFP fluorescence colocalizations (red arrows were visualized in aggregates and autophagic bodies (in the vacuole) in root cells of wild type plants carrying the OEP24.1-GFP and RFP-ATG8e constructs by fluorescence confocal microscopy. A,B,C show representative root images. Scale bars: 10 µm.

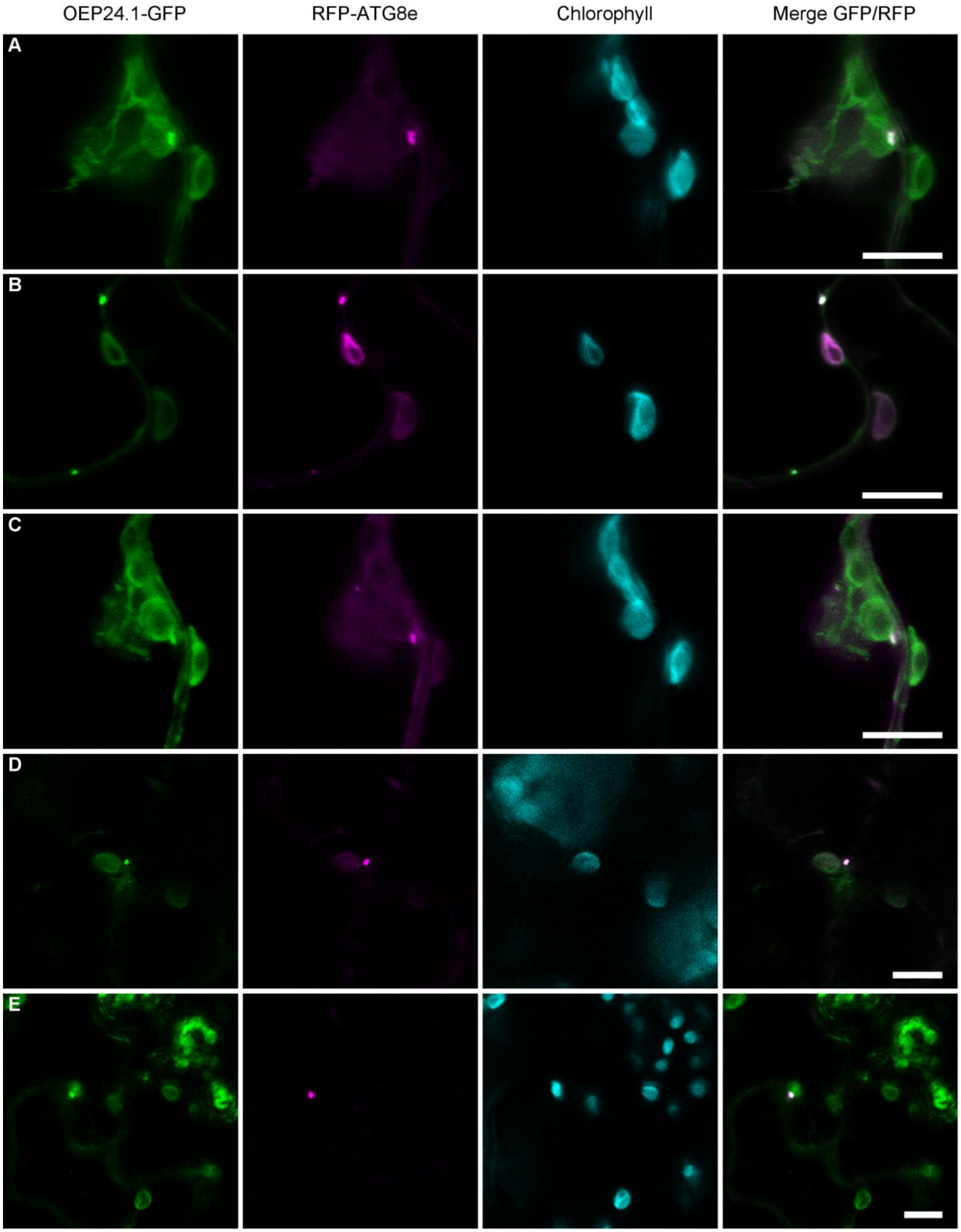

**Supplemental Fig.S11: OEP24.1 is present in autophagosomes in *Nicotiana benthamiana*.** *Nicotiana benthamiana* mesophyll cells transiently expressing the RFP-ATG8e and OEP24.1-GFP constructs after agroinfiltration were analyzed by confocal microscopy. The A-F panels present different cells co-expressing the fusion and representative of different experiments (n=3). In all of them, the colocalization of the GFP and RFP fluorescence show the localization of OEP24.1-GFP in autophagosomes, several of which are close to chloroplast envelopes. Green color shows the fluorescent of the OEP24.1-GFP protein fusion, magenta color shows RFP-ATG8 autophagosome fluorescence, cyan color shows chloroplast autofluorescence. The last image of each panel merges all the signals. Scale bars: 10 µm.

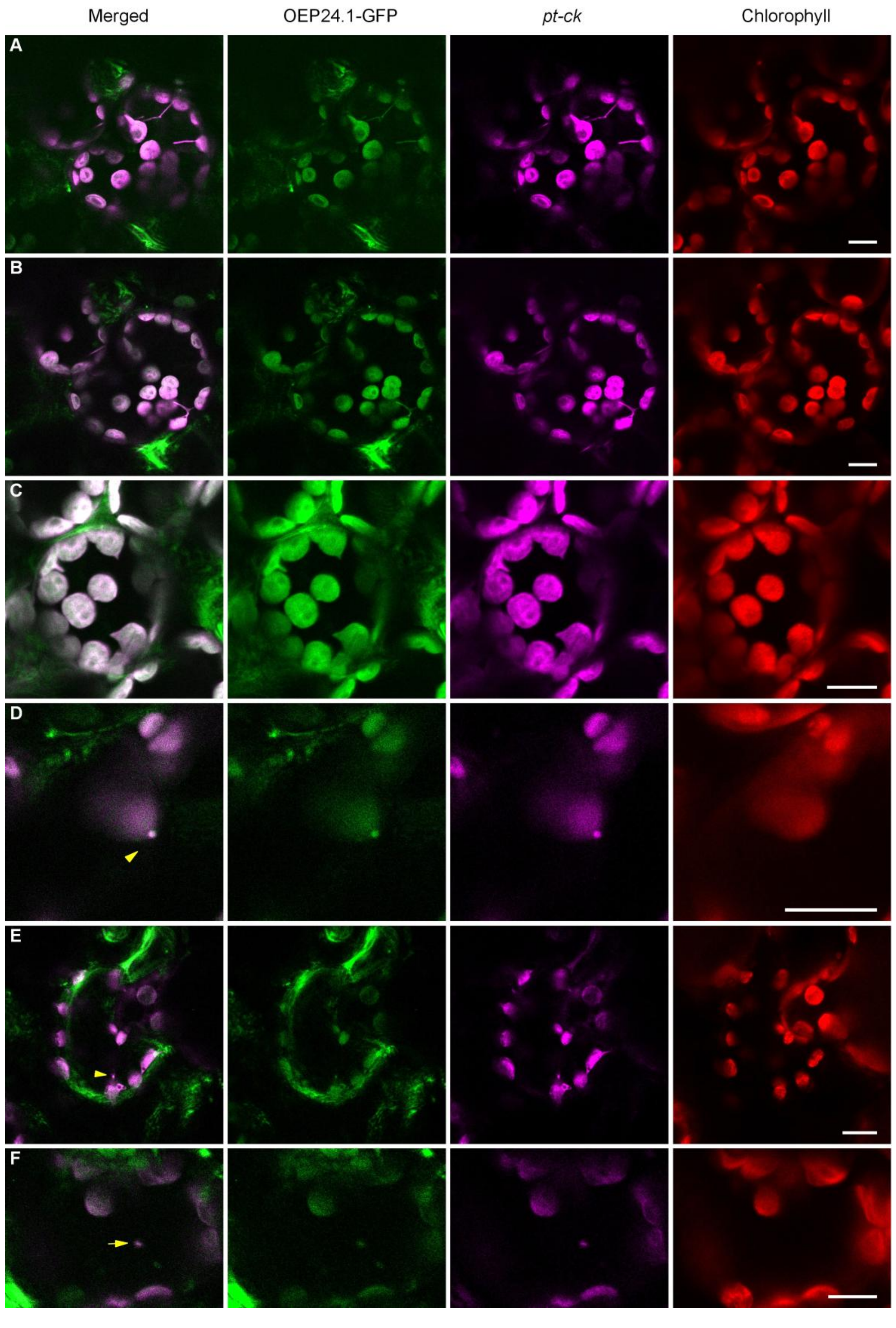

**Supplemental FigS12: Signals of OEP24.1-GFP and *pt-ck* fusions co-colocalize in in chloroplasts and stromules (A,B,C), and in puncta detaching from chloroplast (D) and mobile puncta in the cytosol (E,F).** Mesophyll cells of *Nicotiana benthamiana* transiently expressing and OEP24.1-GFP and *pt-ck* constructs following agroinfiltration were analyzed by confocal microscopy. Panels A-F show images acquired 2 days after agroinfiltration. Green fluorescence corresponds to the OEP24.1–GFP fusion protein, magenta fluorescence corresponds to CFP fused to stromal protein (*pt-ck*), and red fluorescence corresponds to chloroplast autofluorescence. The first image in each panel shows a merge of green and magenta signals and arrows point at co-localisations. Scale bars: 10 µm.

**A. mRNA sequence of Arabidopsis thaliana outer envelope pore 24B-like protein (AT5G42960), fasta code: NM_123662.3**

GCCGCTCTCACCGAACCTCTGCTTTCTCTCTCTCTCTCTCTCTCTCTCTCTCTCTCTCTCTCTC
TCTCTATCTCTACTGTGCGAAAATGGCGATGAAGGCTTCTATCAAAGGCAAGTACGATACTGACAAAACC
AGTGGAATCGGTTCACTTGCTTTCAACGCCGGCGATATTAAGCTACGCGCCACCATGACCGATGCTACTC
TCGTCGCCGGTCCTACTTTGACCGGTCTCGCTCTCGCTGTAGAGAAGCCTGGTTCCTTCATCGTCGAGTA
CAACGTCCCTAAAAAGGATGTTAGGTTTCAGTTCATGAACACAGTAAGAATTGCAGAGAAGCCTTTGAAT
TTGACTTACATTCATAGCAGAGCTGATAACAGAACAATTGTCGATGGGAGCCTTGTGATTGATTCTGCAA
ACAAACTCTCTGCTAACCACATGGTTGGGACAAACAACTGTAAGATTAAGTATACCTATGCTCATGGAGG
GTTAGCTACGTTCGAGCCGTGTTATGACTTGGCGAAAAATACATGGGATTTTGCGGTTTCTCGAAGATTT
TATAGTGGTGATAATGTGAGGGCAACTTATCAGACGTCTAGTAAGTTGTTGGGTATGGAATGGTCAAGGA
ACAACAAAGCAAGTGGATTCAAGGTATGTGCGTCTGTGAATCTTGCGGATGAACTGAAAACGCCGAAATT
GACCGCAGAAACTACATGGAACCTTGAAATGTAGTCTCTTCTTTACACTACTTAGCTAAGTTTCACTTCA
ATTGTTTGGTGTTGCAACACATTTTACTAGTCGGTTCTTGCGAATCTTCCGAAATTTTGTTGCAACAGTT
TCTCATTATTATTACTAGCAGTTTCGTCTGTTATCATTGATCATTTCTTAAATTCGCATAAGTTTTGATC
GTTCAAGAATAAAAATTAAACAACG

**B. Sequence of Arabidopsis thaliana outer envelope pore 24B-like protein**

MAMKASIKGKYDTDKTSGIGSLAFNAGDIKLRATMTDATLVAGPTLTGLALAVEKPGSFIVEYNVPKKDVRFQFMNTVRIAEKPLN
LTYIHSRADNRTIVDGSLVIDSANKLSANHMVGTNNCKIKYTYAHGGLATFEPCYDLAKNTWDFAVSRRFYSGDNVRATYQTSSKL
LGMEWSRNNKASGFKVCASVNLADELKTPKLTAETTWNLEM

**C. Sequence of the cDNA of the OEP24.1 4.3 CRISPR mutant of Arabidopsis thaliana outer envelope pore 24B-like protein (Deletion is indicated by dashed line)**

GCCGCTCTCACCGAACCTCTGCTTTCTCTCTCTCTCTCTCTCTCTCTCTCTCTCTCTCTCTCTC
TCTCTATCTCTACTGTGCGAAAATGGCGATGAAGGCTTCTATCAAAGGCAAGTACGATACTGACAAAACC
AGTGGTGACCGG..........................................................................................................................................
........................................................TCTCGCTCTCGCTGTAGAGAAGCCTGGTTCCTTCATCGTCGAGTA
CAACGTCCCTAAAAAGGATGTTAGGTTTCAGTTCATGAACACAGTAAGAATTGCAGAGAAGCCTTTGAAT
TTGACTTACATTCATAGCAGAGCTGATAACAGAACAATTGTCGATGGGAGCCTTGTGATTGATTCTGCAA
ACAAACTCTCTGCTAACCACATGGTTGGGACAAACAACTGTAAGATTAAGTATACCTATGCTCATGGAGG
GTTAGCTACGTTCGAGCCGTGTTATGACTTGGCGAAAAATACATGGGATTTTGCGGTTTCTCGAAGATTT
TATAGTGGTGATAATGTGAGGGCAACTTATCAGACGTCTAGTAAGTTGTTGGGTATGGAATGGTCAAGGA
ACAACAAAGCAAGTGGATTCAAGGTATGTGCGTCTGTGAATCTTGCGGATGAACTGAAAACGCCGAAATT
GACCGCAGAAACTACATGGAACCTTGAAATGTAGTCTCTTCTTTACACTACTTAGCTAAGTTTCACTTCA
ATTGTTTGGTGTTGCAACACATTTTACTAGTCGGTTCTTGCGAATCTTCCGAAATTTTGTTGCAACAGTT
TCTCATTATTATTACTAGCAGTTTCGTCTGTTATCATTGATCATTTCTTAAATTCGCATAAGTTTTGATC
GTTCAAGAATAAAAATTAAACAACG

**D. Sequence of the OEP24.1 4.3 CRISPR mutant truncated protein (Grey sequence indicate the result of the change of translation phase).**

MAMKASIKGKYDTDKTSGDRSRSRCREAWFLHRRVQRP-

**E. Sequence of the cDNA of the OEP24.1 8.2 CRISPR mutant of Arabidopsis thaliana outer envelope pore 24B-like protein (INDEL is indicated in red)**

GCCGCTCTCACCGAACCTCTGCTTTCTCTCTCTCTCTCTCTCTCTCTCTCTCTCTCTCTCTCTC
TCTCTATCTCTACTGTGCGAAAATGGCGATGAAGGCTTCTATCAAAGGCAAGTACGATACTGACAAAACC
AGTGGAAAGTAGGACCGGCGACGAGAGTAGCATCGGTCATGGTGGCGCGTAGCTTAATATCGCCGGCGTT
GAAAGCAAGTGAACCGATTGACCGGTCTCGCTCTCGCTGTAGAGAAGCCTGGTTCCTTCATCGTCGAGTA
CAACGTCCCTAAAAAGGATGTTAGGTTTCAGTTCATGAACACAGTAAGAATTGCAGAGAAGCCTTTGAAT
TTGACTTACATTCATAGCAGAGCTGATAACAGAACAATTGTCGATGGGAGCCTTGTGATTGATTCTGCAA
ACAAACTCTCTGCTAACCACATGGTTGGGACAAACAACTGTAAGATTAAGTATACCTATGCTCATGGAGG
GTTAGCTACGTTCGAGCCGTGTTATGACTTGGCGAAAAATACATGGGATTTTGCGGTTTCTCGAAGATTT
TATAGTGGTGATAATGTGAGGGCAACTTATCAGACGTCTAGTAAGTTGTTGGGTATGGAATGGTCAAGGA
ACAACAAAGCAAGTGGATTCAAGGTATGTGCGTCTGTGAATCTTGCGGATGAACTGAAAACGCCGAAATT
GACCGCAGAAACTACATGGAACCTTGAAATGTAGTCTCTTCTTTACACTACTTAGCTAAGTTTCACTTCA
ATTGTTTGGTGTTGCAACACATTTTACTAGTCGGTTCTTGCGAATCTTCCGAAATTTTGTTGCAACAGTT
TCTCATTATTATTACTAGCAGTTTCGTCTGTTATCATTGATCATTTCTTAAATTCGCATAAGTTTTGATC
GTTCAAGAATAAAAATTAAACAACG

**F. Sequence of the OEP24.1 8.2 CRISPR mutant truncated protein**

MKASIKGKYDTDKTSGK-

**Supplemental Fig.S13: OEP24.1 wild type and mutated cDNA sequences (A,C,E) and protein sequences (B,D,F).** The cDNA and protein sequences of wild type (A,B), #4.2 CRISPR mutant (C,D) and #8.2 CRISPR mutant (E,F) are presented. On the cDNA sequences, guide RNAs used for mutation are tagged in green with their CGG sequences tagged in red. The ATG and TAG start and stop codons of OEP24.1 are in red. New STOP codons introduced by mutations are tagged in yellow and new sequences introduced by deletion in #8.2 is in red. On protein sequences, sequence modifications of the #4.2 and #8.2 OEP24.1 proteins are in grey.

A

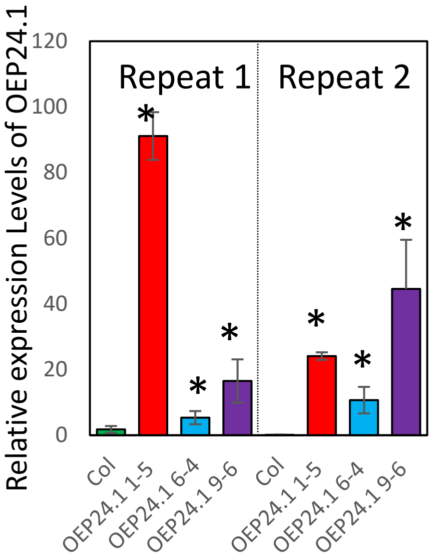

B

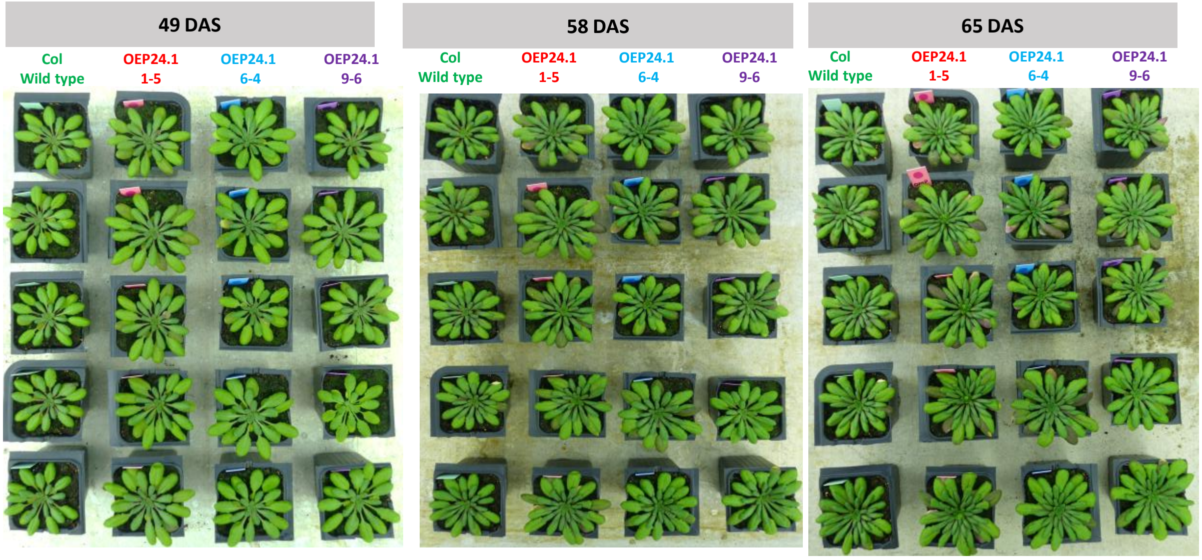

**Fig.S14: *OEP24.1* overexpressing lines do not display any different phenotype from wild type. (**A) Relative expression level of OEP24.1 gene in wild type and *OEP24.1* overexpressing lines. Expression level was monitored using qRT-PCR using primers described in Table S2 and their relative units (Y axis) were obtained normalizing to a synthetic reference gene that combines EF1-$\alpha$ and APC2. Mean and SD are shown (n=4) and significant differences between overexpressing lines and WT are indicated by * (P<0.5). (B) Phenotypes of the rosettes of wild type and *OEP24.1* overexpressing plants grown under low nitrate conditions under short days. The same four plants are shown at 49, 58 and 65 days after sowing. Culture run was repeated three times, under the same condition with similar results.

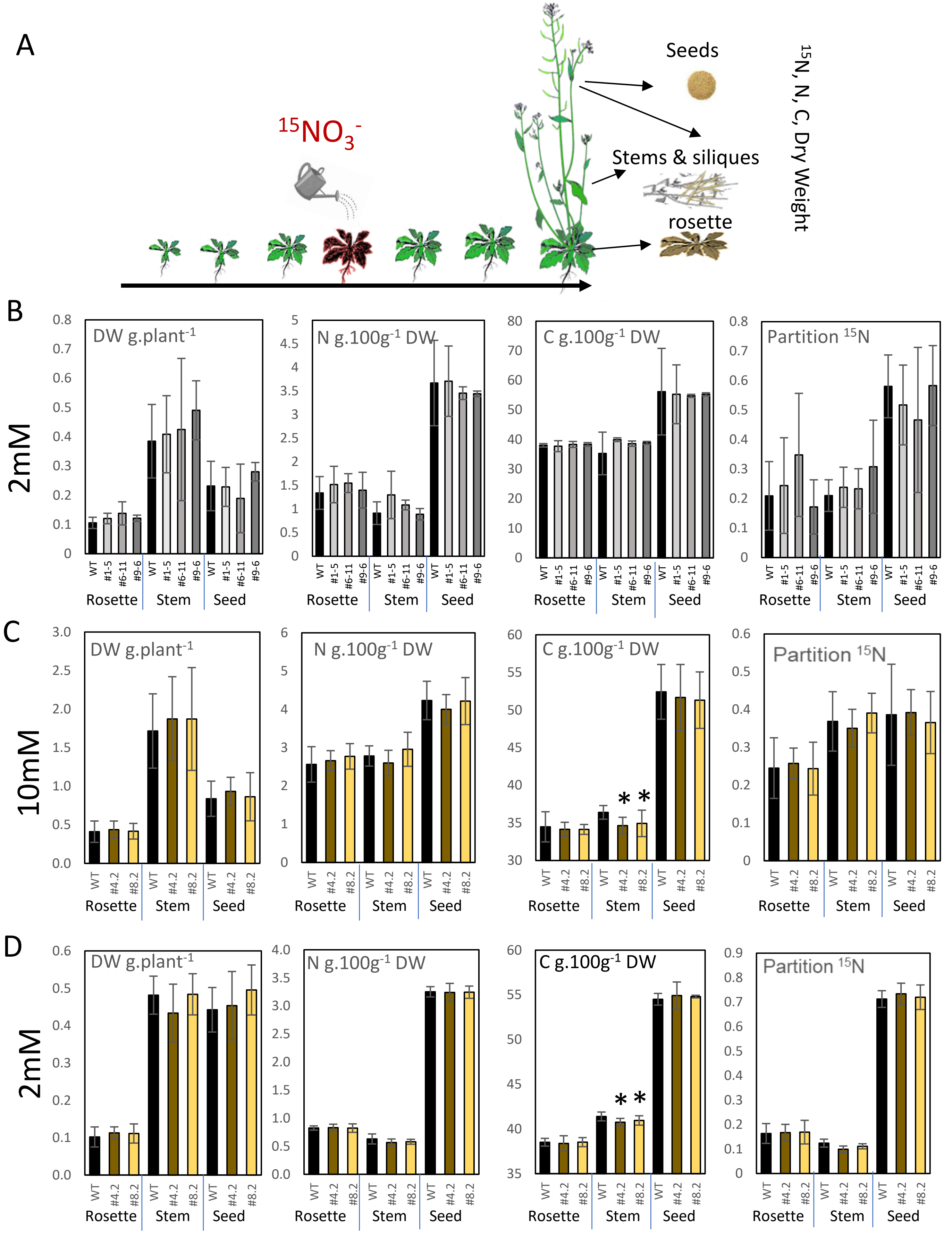

**Supplemental Fig.S15: Carbon concentration is lower in the stem of OEP24.1 CRISPR mutants. (A)** Schematic representation of the culture experimental design of $^{15}$N tracing experiment and organ harvest at plant maturity for the determination of $^{15}$N, N and C concentrations and organ biomasses (see material and methods). Plant biomass (DW), N concentrations and C concentrations were measured in the different organs (rosette, stem and siliques, and seeds) of OEP24.1 overexpressing lines (B), *oep24.1* CRISPR mutants grown under high nitrate supply (C), or un low nitrate supply (D). $^{15}$N partitioning was calculated as a percentage dividing the $^{15}$N content (mg) in the different plant organs by the total $^{15}$N content (mg) in the whole plant. Mean and standard deviation are shown (n=5 for overexpressing lines and n=12 for CRISPR mutants). Significant difference between wild type and mutants or overexpressing lines are shown by * (P<0.05).

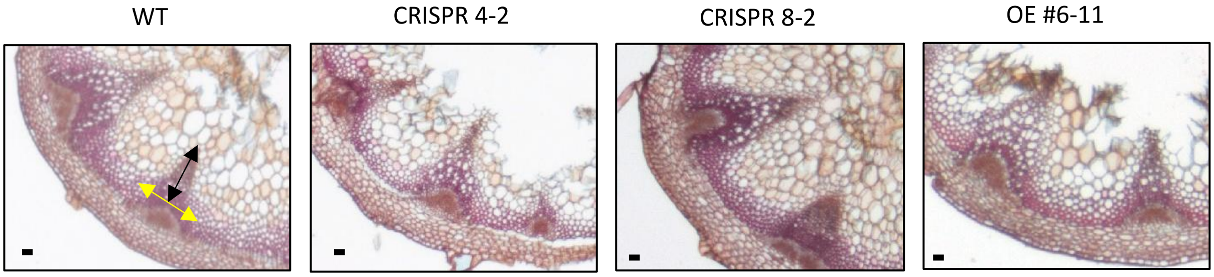

**Supplemental Fig.S16:** Representative transverse stem sections of wild type (WT), CRISPR mutants (4-2 and 8-2), and OE #6-11 over-expressing line. Stem sections were stained using Alcian Blue/Safranin (2:1). Yellow and Dark arrows indicate xylem length and width respectively. Scale bar = 50 µm

**Supplemental Table S1:** All the plasmids listed in this table have been used in this study.

| Plasmid | Description | Reference |
|---|---|---|
| **Gateway cloning vectors** | | |
| pDONR207 | Gateway® Donor Vector with attP1 and attP2 sites, $Genta^R$ | Invitrogen |
| pMDC32 | Gateway® Compatible Plant Transformation Vector, $Hyg^R$, $Kan^R$ | ABRC |
| pUBN-Dest-eGFP | Gateway® Compatible Plant Transformation Vector Ubiquitin10 promoter, eGFP fusion in C terminal, $Basta^R$ $Spec^R$ | Grefen et al. 2010 |
| pUBN-Dest-cYFP | Gateway® Compatible Plant Transformation Vector, Ubiquitin10 promoter, cYFP fusion in C terminal $Basta^R$ $Spec^R$ | Grefen et al. 2010 |
| pUBN-Dest-nYFP | Gateway® Compatible Plant Transformation Vector, Ubiquitin10 promoter, nYFP fusion in C terminal $Basta^R$ $Spec^R$ | Grefen et al. 2010 |
| pH7WGR2 | Gateway® Compatible Plant Transformation Vector, p35S promotor, RFP fusion in N terminal, $Hygro^R$, $Spec^R$ | VIB-UGENT center for pant system biology |
| pGADT7 | Gateway® Destination Vector for yeast two hybrid, $Amp^R$ | Deruyffelaere et al. 2018 |
| pGBKT7 | Gateway® Destination Vector for yeast two hybrid, $Kana^R$ | Deruyffelaere et al. 2018 |
| **pUNI51, pENTR223, pDONR207 cloning vector** | | |
| U15871 | pENTR/SD-dTopo Cloning Vector with OEP24.1 ORF | ABRC |
| pDONR207-ATG8a | pDONR207 with ATG8a coding DNA plus introns (from ATG to STOP) | Chen et al.2019 |
| G22921 | pENTR223Gateway TM entry vector with ATG8b ORF | ABRC |
| G51304 | pENTR223Gateway TM entry vector with ATG8c ORF | ABRC |
| G22544 | pENTR223Gateway TM entry vector with ATG8d ORF | ABRC |
| pDONR207-ATG8e | pDONR207 Donor Vector with ATG8d ORF | Merkulova et al. 2014 |
| G21342 | pENTR223Gateway TM entry vector with ATG8f ORF | ABRC |
| U17226 | PENTR/SD-DTOPOGateway TM entry vector with ATG8g ORF | ABRC |
| G82070 | pENTR223Gateway TM entry vector with ATG8h ORF | ABRC |
| G82060 | pENTR223Gateway TM entry vector with ATG8i ORF | ABRC |
| **Entry clones made by subcloning** | | |
| pDONR207-OEP24.1 STOP less | Gateway® pDONR207 vector with Arabidopsis OEP24.1 CDS STOP-less, $Genta^R$ | This study |
| **Entry clones for site directed mutagenesis** | | |
| pDONR221-ATG8d ΔLDS | Gateway® pDONR207 vector with ATG8d CDS lacking LDS, $Kan^R$ | This study |
| pDONR221-ATG8e ΔLDS | Gateway® pDONR221 vector with ATG8e CDS lacking LDS, $Kan^R$ | This study |
| pDONR207-ATG8e ΔUDS | Gateway® pDONR207 vector with Arabidopsis ATG8e lacking UDS, $Genta^R$ | This study |
| pENTR223-ATG8d ΔUDS | Gateway® pENTR223 vector with Arabidopsis ATG8d lacking UDS, $Genta^R$ | This study |
| **Plant expression vectors made by subcloning or ordered** | | |

| Plasmid | Description | Reference |
|---|---|---|
| p35S::RFP-ATG8e | ATG8e CDS in pH7WGR2, $Hygro^R$ | This study |
| pUBi::OEP24.1-GFP | OEP24.1 CDS in pUBC-Dest-eGFP, $Basta^R$ | This study |
| p35S::OEP24.1 | OEP24.1 CDS in PMDC32. $Hygro^R$ | This study |
| pUBi::cYFP-OEP24.1 | OEP24.1 CDS in pUBN-Dest-cYFP, $Basta^R$ | This study |
| pUBi::nYFP-OEP24.1 | OEP24.1 CDS in pUBN-Dest-nYFP, $Basta^R$ | This study |
| pUBi::cYFP-ATG8g | ATG8g CDS in pUBN-Dest-cYFP, $Basta^R$ | This study |
| pUBi::nYFP-ATG8g | ATG8g CDS in pUBN-Dest-nYFP, $Basta^R$ | This study |
| pUBi::cYFP-ATG8d | ATG8d CDS in pUBN-Dest-cYFP, $Basta^R$ | This study |
| pUBi::nYFP-ATG8d | ATG8d CDS in pUBN-Dest-nYFP, $Basta^R$ | This study |
| PRMCas9 OEP24 | Crispr Cas 9 vector including guide RNAs targeting OEP24.1 $Kana^R$,$Hygro^R$, | This study |
| pt-yk | CD3-997 ABRC stock | Nelson et al. 2007 |
| pt-ck | CD3-993 ABRC stock | Nelson et al. 2007 |
| **Yeast expression vectors made by subcloning** | | |
| pGADT7-OEP24.1 | pGAD derived vector with OEP24.1 fused to the Gal4 AD in N terminal; for yeast two hybrid, $Ampi^R$ | This study |
| pGBKT7-ATG8a | pGBK derived vector with ATG8a fused to the Gal4 BD in N terminal; for yeast two hybrid, $Kan^R$ | This study |
| pGBKT7-ATG8b | pGBK derived vector with ATG8b fused to the Gal4 BD in N terminal; for yeast two hybrid, $Kan^R$ | This study |
| pGBKT7-ATG8c | pGBK derived vector with ATG8c fused to the Gal4 BD in N terminal; for yeast two hybrid, $Kan^R$ | This study |
| pGBKT7-ATG8d | pGBK derived vector with ATG8d fused to the Gal4 BD in N terminal; for yeast two hybrid, $Kan^R$ | This study |
| pGBKT7-ATG8e | pGBK derived vector with ATG8e fused to the Gal4 BD in N terminal; for yeast two hybrid, $Kan^R$ | This study |
| pGBKT7-ATG8f | pGBK derived vector with ATG8f fused to the Gal4 BD in N terminal; for yeast two hybrid, $Kan^R$ | This study |
| pGBKT7-ATG8g | pGBK derived vector with ATG8g fused to the Gal4 BD in N terminal; for yeast two hybrid, $Kan^R$ | This study |
| pGBKT7-ATG8h | pGBK derived vector with ATG8h fused to the Gal4 BD in N terminal; for yeast two hybrid, $Kan^R$ | This study |
| pGBKT7-ATG8i | pGBK derived vector with ATG8i fused to the Gal4 BD in N terminal; for yeast two hybrid, $Kan^R$ | This study |
| pGBKT7-ATG8e ΔLDS | pGBK derived vector with ATG8e lacking LDS site fused to the Gal4 BD in N terminal; for yeast two hybrid, $Kan^R$ | This study |
| pGBKT7-ATG8d ΔLDS | pGBK derived vector with ATG8d lacking LDS site fused to the Gal4 BD in N terminal; for yeast two hybrid, $Kan^R$ | This study |
| pGBKT7-ATG8e ΔUDS | pGBK derived vector with ATG8e lacking UDS site fused to the Gal4 BD in N terminal; for yeast two hybrid, $Kan^R$ | This study |
| pGBKT7-ATG8d ΔUDS | pGBK derived vector with ATG8d lacking UDS site fused to the Gal4 BD in N terminal; for yeast two hybrid, $Kan^R$ | This study |

Supplemental Table S2: Primers used in this study.

| GENE | NAME | | FORWARD | REVERSE |
|---|---|---|---|---|
| gateway | U3-U5 | Cloning | GGGGACCACTTTGTACAAGAAAGCTGGGTCTCCACCTCCGGATC | GGGGACAAGTTTGTACAAAAAAGCAGGCTTCGAAGGAGATAGAACCATG |
| AT5G42960 | gtwOEP24 | Cloning | GGAGATAGAACCATGGCGATGAAGGCTTCTATC | TCCACCTCCGGATCCCATTTCAAGGTTCCATGTAGTTTC |
| AT2G45170 | gtwATG8e | Cloning | GGAGATAGAACCATGAATAAAGGAAGCATC | TCCACCTCCGGATCAGATTGAAGAAGCACCGAATG |
| AT5G42960 | crisprOEP | PCR | CTACTGTGCGAAAATGGCGA | ATCACAAGGCTCCCATCGAC |
| AT5G42960 | q-OEP24 | RT-qPCR | TTCACTTGCTTTCAACGCCG | ATCACAAGGCTCCCATCGAC |
| AT5G60390 | q-EF-1α | RT-qPCR | CTGGAGGTTTTGAGGCTGGTAT | CCAAGGGTGAAAGCAAGAAGA |
| AT2G04660 | q-APC2 | RT-qPCR | GAAACATCAATTGCCTCTGTGGAAGA | AAGGATCAGCCACACAAAACATCTTG |
